# BBLN-1 is essential for intermediate filament organization and apical membrane morphology

**DOI:** 10.1101/2020.12.04.411520

**Authors:** Sanne Remmelzwaal, Florian Geisler, Riccardo Stucchi, Suzanne van der Horst, Milena Pasolli, Jason R. Kroll, Olga D. Jarosinska, Anna Akhmanova, Christine A. Richardson, Maarten Altelaar, Rudolf E. Leube, João J. Ramalho, Mike Boxem

## Abstract

Epithelial tubes are essential components of metazoan organ systems that control the flow of fluids and the exchange of materials between body compartments and the outside environment. The size and shape of the central lumen confer important characteristics to tubular organs and need to be carefully controlled. Here, we identify the small coiled-coil protein BBLN-1 as a regulator of lumen morphology in the *C. elegans* intestine. Loss of BBLN-1 causes the formation of bubble-shaped invaginations of the apical membrane into the cytoplasm of intestinal cells, and abnormal aggregation of the subapical intermediate filament (IF) network. BBLN-1 interacts with IF proteins and localizes to the IF network in an IF-dependent manner. The appearance of invaginations is a result of the abnormal IF aggregation, indicating a direct role for the IF network in maintaining lumen homeostasis. Finally, we identify bublin (BBLN) as the mammalian ortholog of BBLN-1. When expressed in the *C. elegans* intestine, bublin recapitulates the localization pattern of BBLN-1 and can compensate for the loss of BBLN-1. In mouse intestinal organoids, bublin localizes subapically, together with the IF protein keratin 8. Our results therefore may have implications for understanding the role of IFs in regulating epithelial tube morphology in mammals.

**Summary:** We identify BBLN-1 as an evolutionary conserved regulator of lumen morphology in the *C. elegans* intestine. Loss of *bbln-1* causes intermediate filament network reorganization that induces severe apical morphology defects. We also identify bublin (BBLN) as the mammalian ortholog, which can compensate for the loss of BBLN-1 in *C. elegans*.

## Introduction

Epithelial and endothelial tubes are fundamental building units of many organs, including the digestive tract, vascular system, kidney, and lung. Tubular organs are essential for the transport of nutrients, waste products, gases, and ions across large distances. Furthermore, epithelial tubes function as a protective barrier to the outside environment. Tubes consist of a central lumen bounded by the apical domains of one or more epithelial or endothelial cells, and are remarkably varied in size, complexity and mechanism of development (Iruela-Arispe and Beitel, 2013; Lubarsky and Krasnow, 2003; Sigurbjörnsdóttir et al., 2014). The size and shape of the central lumen are precisely controlled, as they confer important biophysical and biochemical properties to biological tubes. Lumen morphology can be very stable, but can also be highly plastic during tissue growth or when subjected to mechanical stimuli or stress (Iruela-Arispe and Davis, 2009; Stutz et al., 2015; Sundaram and Buechner, 2016). The importance of mechanisms that control lumen morphology is highlighted by common pathologies that are characterized by altered lumen architecture, including polycystic kidney disease, cystic fibrosis, and inflammatory bowel disease (Bergmann et al., 2018; Cutting, 2015; Ramos and Papadakis, 2019; Schelling, 2016). The *Caenorhabditis elegans* intestine provides a simple model to study the regulation of lumen morphology. The *C. elegans* intestine is a tube composed of 20 cells arranged in nine segments surrounding a central lumen (Fig. 1A) (Sulston et al., 1983; Leung et al., 1999). Specialized cell-cell junctions at the lateral membrane, known as *C. elegans* apical junctions (CeAJ), keep neighboring cells tightly adherent, ensuring integrity of the epithelium and impermeability of the lumen (Pasti and Labouesse, 2014). The intestinal cells are born during embryogenesis and do not divide or renew during larval development or in adulthood. They are polarized along an apicobasal axis and resemble mammalian enterocytes at the ultrastructural level (Coch and Leube, 2016). The lumen forming apical surface of both cell types is covered by microvilli that contain bundled actin filaments, and are supported by the subapical actin-rich terminal web (Leung et al., 1999; Bossinger et al., 2004). The luminal surface is further supported by an intermediate-filament (IF) rich fibrous sheet that directly underlies — and is likely connected to — the terminal web. The *C. elegans* IF network forms a particularly electron-dense structure anchored at the lateral cell junctions that is known as the ‘endotube’ (Bossinger et al., 2004; Coch and Leube, 2016; Munn and Greenwood, 1984).

**Figure 1.**
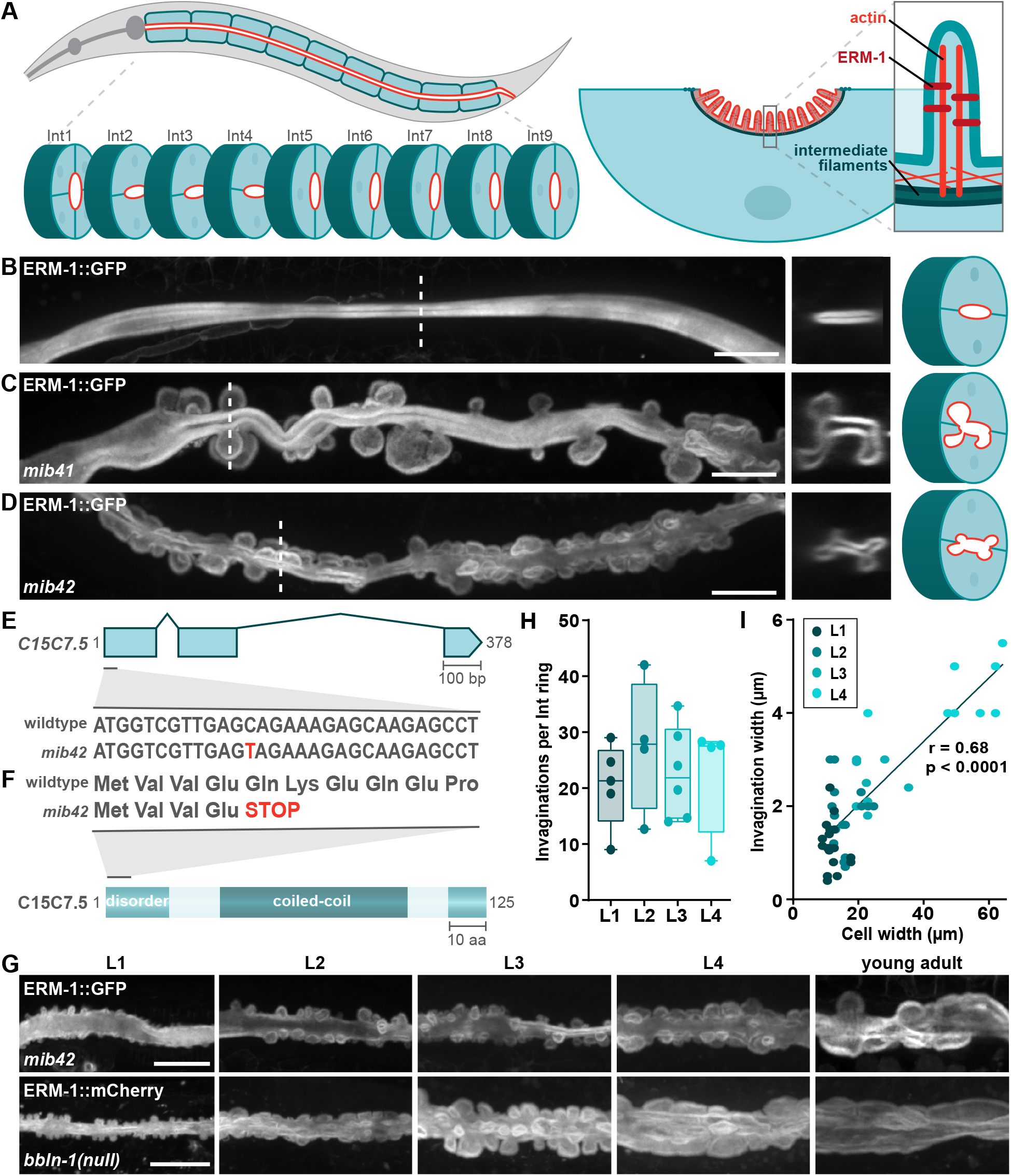
The intestines of *mib41* and *mib42* mutants show cytoplasmic invaginations. (**A**) Schematic representation of the *C. elegans* intestine (left) and the subcellular intestinal localization of proteins and cytoskeletal elements relevant for this study (right). (**B–D**) Intestinal apical membrane morphology in L4 animals visualized with an endogenous ERM-1::GFP reporter. Dotted lines in left panels indicate position of cross-section views. Schematics depict cross-section with apical membrane in red. Unless otherwise indicated, in this and all other figures, images were acquired using spinning-disk confocal microscopy, lateral views are maximum intensity projections, anterior is to the left, and scale bars indicate 10 µm. (**E, F**) Genomic structure of the *C15C7*.*5* gene (E) and representation of the C15C7.5 protein with predicted domains (F). Boxes in the genomic structure correspond to exons while lines represent introns. Part of the DNA and amino acid sequence is shown for both the wild-type and *mib42* mutant allele. (**G**) Progression of *mib42* and *bbln-1*(*null*) invagination phenotype throughout development visualized by endogenous ERM-1::GFP and ERM-1::mCherry reporters, respectively. (**H**) Quantification of number of invaginations per intestinal ring, per larval stage in *bbln-1*(*null*) animals. Each data point represents the average number of invaginations per intestinal ring in one animal. Data is presented as median ± interquartile range. Differences are not significant (Kruskal-Wallis followed by Dunn’s multiple comparisons test; P > 0.99). (**I**) Correlation between cell and invagination width in *bbln-1*(*null*) animals, color coded per larval stage. Linear regression is shown for all data points combined. Data is analyzed with nonparametric Spearman correlation; r = 0.6824, P < 0.0001.

Cytoplasmic IF proteins form resilient and flexible networks that mediate the mechanical properties of cells and tissues and contribute to a diverse range of processes including organelle positioning, intracellular trafficking, and cell motility (Coulombe and Wong, 2004; Etienne-Manneville, 2018). In tubular epithelia, subapical IF networks are ideally positioned to contribute to the regulation of lumen morphology and functioning. Indeed, intestinal IF networks have essential roles in epithelial polarization, maintenance of lumen morphology, and protection from environmental stresses (Etienne-Manneville, 2018; Geisler and Leube, 2016; Salas et al., 2016; Sanghvi-Shah and Weber, 2017; Toivola et al., 2010). In the *C. elegans* intestine, loss of IF subunits, or of the intermediate filament organizers IFO-1 and SMA-5, can result in morphological abnormalities of the lumen and increased susceptibility to microbial, osmotic, and oxidative stresses (Carberry et al., 2012; Geisler et al., 2016, 2019, 2020).

Here, we report the identification of an evolutionary conserved regulator of intestinal tube morphology and IF organization we term *bbln-1*, for <u>b</u>ulges <u>b</u>udding from the lume<u>n</u>. *bbln-1* mutants are characterized by the presence of large invaginations of the apical lumen into the cytoplasm of intestinal cells. The BBLN-1 protein physically interacts with IF proteins and localizes to the endotube in an IF-dependent manner. Loss of BBLN-1 causes defects in the organization of the intestinal IF network, which aggregates into thick bundles at the neck of cytoplasmic invaginations. Surprisingly, the invagination phenotype of *bbln-1* mutants is rescued by removal of the IF network. This indicates that the cytoplasmic invaginations are the result of the aggregated IF network state caused by loss of *bbln-1*. Based on sequence similarity, we identified a mammalian ortholog of BBLN-1, which we termed bublin (BBLN). Bublin shares multiple protein interaction partners with BBLN-1, and can partially substitute for BBLN-1 functioning in the *C. elegans* intestine. Our findings provide further evidence that the subapical IF network plays an active role in regulating the morphology of tubular epithelia, while the structural and functional conservation from *C. elegans* to humans provides a model for studying IF modulation *in vivo*.

## Results

### Loss of bbln-1 causes cytoplasmic invaginations of the intestinal apical membrane

To investigate the mechanisms that regulate intestinal lumen morphology, we performed a genetic screen in *C. elegans* for mutants with lumen morphology defects. We mutagenized a strain expressing the apical membrane protein ERM-1 endogenously fused to GFP, and screened for viable F2 progeny with aberrations in lumen morphology (Fig. 1A–D). We identified two independent mutants (*mib41* and *mib42*) with dramatic bubble-shaped protrusions of the apical plasma membrane into the cytoplasm of the intestinal cells (Fig. 1C, D). Complementation analysis indicated that *mib41* and *mib42* affect two independent loci. We determined the genomic regions of interest by single nucleotide polymorphism (SNP) mapping (Davis et al., 2005), and identified candidate causative mutations using the sibling subtraction method for mapping by whole-genome sequencing (Joseph et al., 2018). We obtained a single cluster of candidate mutations for each mutant allele on opposite arms of the X chromosome, consistent with the SNP mapping data.

Only one among the candidate *mib41* mutations was predicted to exert a detrimental effect on protein function, by inducing a missense mutation within the *sma-5* open reading frame (ORF) (C370T in *sma-5a*). At the protein level, this mutation results in an arginine to tryptophan substitution in the kinase domain of all predicted isoforms (Arg124Trp in SMA-5a) (Fig. S1A). SMA-5 is a member of the conserved mitogen-activated protein kinase (MAPK) family that is required for intestinal lumen morphogenesis (Geisler et al., 2016). *sma-5* mutations induce a lumen invagination phenotype similar to *mib41* mutants (Geisler et al., 2016) and the strong loss of function *sma-5*(*n678*) allele (Watanabe et al., 2005) failed to complement *mib41*. Together, these data show that *mib41* is a novel *sma-5* allele.

The *mib42* sequencing data also yielded a single potentially deleterious mutation within a predicted ORF: a C to T transition resulting in an early stop at the fifth codon of the uncharacterized gene *C15C7*.*5* (Fig. 1E, F). We used several strategies to confirm that loss of *C15C7*.*5* function was responsible for the intestinal lumen morphology defects in *mib42* animals. First, *C15C7*.*5* RNA interference (RNAi) feeding experiments in ERM-1::GFP animals revealed apical membrane invaginations similar to those observed in the mutant (Fig. S1B). Second, transgenic expression of *GFP::C15C7*.*5* from the intestinal specific *vha-6* promoter fully rescued the invagination phenotype of *mib42* animals (Fig. S1C). Lastly, we generated a *C15C7*.*5* null allele, *mib70*, by removing the entire *C15C7*.*5* coding sequence using CRISPR/Cas9 engineering. The *mib70* deletion, from here on referred to as *C15C7*.*5*(*null*), caused cytoplasmic invaginations (Fig. S1D) and was unable to complement *mib42*. Both *C15C7*.*5*(*mib42*) and *C15C7*.*5*(*null*) animals are homozygous viable, but animals appear somewhat sick and display a slight clear phenotype. The invagination phenotype observed in *C15C7*.*5*(*null*) animals was more severe than in *mib42* mutants (Fig. 1G). This data indicates that *mib42* is a partial loss-of-function mutation in the *C15C7*.*5* gene, and that C15C7.5 controls intestinal lumen morphology. We have named the *C15C7*.*5* gene *bbln-1* for <u>b</u>ulges <u>b</u>udding from the intestinal lume<u>n</u>.

We next examined the intestinal cytoplasmic invaginations in more detail. In *bbln-1*(*null*) animals, invaginations were visible from hatching onwards, and the number of invaginations per intestinal ring remained constant throughout development (Fig. 1G, H). Invagination width correlated with intestinal cell width, and in late larval and adult stages invaginations had an elongated appearance (Fig. 1I). The *bbln-1* invaginations were spread uniformly along the length of the intestine (Fig. 1D, G). However, in cross section, invaginations seemed to preferentially localize at the vertices of the ellipse shaped lumen, in proximity to cell junctions. This was particularly noticeable in *mib42* animals, which have fewer invaginations than the *bbln-1*(*null*) mutant (Fig. 1D). To confirm this, we visualized invaginations in a *bbln-1*(*mib42*) strain expressing the junctional protein DLG-1 endogenously tagged with mCherry. Indeed, invaginations occurred in proximity to cell junctions (Fig. S1E–H).

Finally, we assessed whether invaginations are contiguous with the lumen by feeding animals with a membrane impermeable Texas Red-dextran conjugate. In wild-type animals, Texas Red signal filled the lumen and was enclosed by the ERM-1::GFP-labelled apical membrane (Fig. S1I). Similarly, in *bbln-1*(*mib42*) animals, all invaginations contained Texas Red signal, indicating that they are contiguous with the lumen. Moreover, we never observed internalized invaginations or presence of Texas Red signal in the cytoplasm of intestinal cells, indicating that the invaginations are stable and do not internalize.

### BBLN-1 is a small coiled-coil protein that localizes dynamically to intermediate filament-rich structures

*bbln-1* is predicted to encode a single protein isoform of just 125 amino acids. A central coiled-coil domain (aa 39–102) is the only recognizable feature, with the remainder of the protein predicted to be intrinsically disordered (Fig. 1F). To investigate the function of BBLN-1 in intestinal lumen morphogenesis, we first analyzed its expression pattern and subcellular localization. We used CRISPR/Cas9 engineering to introduce the GFP coding sequence immediately upstream of the *bbln-1* start codon. Homozygous knock-in animals expressing GFP::BBLN-1 do not show obvious developmental or intestinal lumen defects, demonstrating that the fusion protein is functional. We also generated a C-terminally mKate2 tagged variant. However, homozygous *bbln-1::mKate2* animals displayed the intestinal cytoplasmic invagination phenotype, indicating a non-functional protein fusion. We therefore used the N-terminal GFP fusion for our experiments.

BBLN-1 was first apparent at the comma stage of embryonic development, at the apical domain of intestinal cells (Fig. 2A). In subsequent embryonic stages, BBLN-1 became visible in the hypodermis and pharynx as well. During larval and adult stages, we detected BBLN-1 in many epithelial tissues including the pharynx, intestine, excretory canal, hypodermis, vulva, and uterine seam (Fig. 2B). In each of these tissues the distribution of BBLN-1 was similar to previously reported distribution of intermediate filament proteins. In the marginal cells of the pharynx, we observed a pattern of short radial filaments, reminiscent of the IF containing tonofilaments (Fig. 2Bi) (Karabinos et al., 2003). In the excretory canal, BBLN-1 localization was similar to that of the intermediate filament proteins IFA-4, IFB-1, and EXC-2/IFC-2 (Fig. 2Bii) (Al-Hashimi et al., 2018; Khan et al., 2019). In the hypodermis, we observed a pattern strongly resembling that of intermediate filaments in the *C. elegans* hemidesmosomes (Fig. 2Biii) (Pasti and Labouesse, 2014; Woo et al., 2004). We also observed expression of BBLN-1 in the uterine seam and vulva (Fig. 2Biv), as previously reported for IFA-1, MUA-6/IFA-2, IFB-1 and EXC-2/IFC-2 (Al-Hashimi et al., 2018; Hapiak et al., 2003; Karabinos et al., 2003; Williams et al., 2015; Woo et al., 2004).

**Figure 2.**
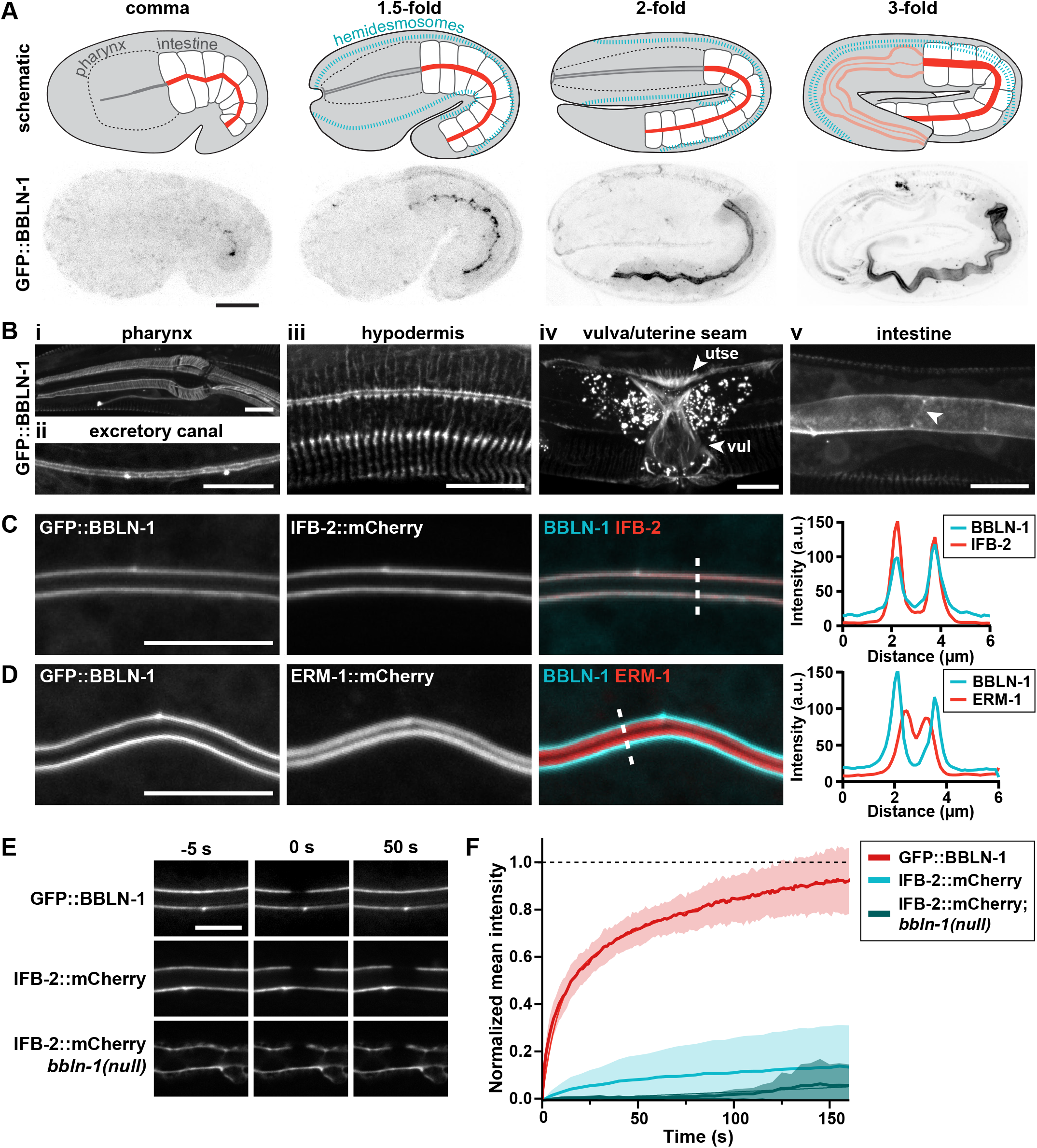
BBLN-1 localizes dynamically to intermediate filament-rich structures. (**A, B**) Distribution of the endogenous GFP::BBLN-1 reporter during late embryonic development (A) and in different tissues in L3 (intestine) and L4 larvae (pharynx and excretory canal images), and young adults (hypodermis and vulva images) (B). (**C, D**) Distribution of GFP::BBLN-1 compared with ERM-1::mCherry (C) and IFB-2::mCherry (D) at the apical domain of intestinal cells of L3 larvae. Dashed lines indicate sites of intensity profiles shown in the graphs to the right. (**E, F**) Stills from time-lapse movies (E) and FRAP curves (F) corresponding to bleached portions of the apical domain in larval intestines. Time-lapse data was acquired at 1 s intervals for GFP::BBLN-1 (n = 7) and 5 s intervals for IFB-2::mCherry (WT: n = 6, *bbln-1*(*null*): n = 5). Thin lines and shading represent the mean ± SD. GFP::BBLN-1 plateau = 0.995, fast half-time = 5.45 s, slow half-time = 55.72, R^2^ = 0.998; IFB-2::mCherry plateau = 0.188, fast half-time = 8.8 s, slow half-time = 97.32, R^2^ = 0.998. For IFB-2::mCherry in a *bbln-1*(*null*) background, R^2^ = 0.817 and no plateau or half-times could be accurately determined.

In the intestine, BBLN-1 localized subapically with apparent accumulation around cell junctions (Fig. 2Bv). This localization pattern corresponds to the distribution of the endotube (Bossinger et al., 2004). To confirm that BBLN-1 resides at the endotube, we generated an endogenous mCherry fusion of the intestinal intermediate filament protein IFB-2. We then analyzed the localization of BBLN-1 relative to IFB-2 and to ERM-1, which localizes apically at microvilli (Bidaud-Meynard et al., 2019; Ramalho et al., 2020). Consistent with localization to the endotube, we found that BBLN-1 colocalizes with IFB-2, basal to ERM-1 (Fig. 2C, D). To determine if BBLN-1 stably associates with the IF network, we performed fluorescence after photobleaching (FRAP) experiments. IFB-2 fluorescence recovered only ∼15% in 150 seconds (Fig. 2E, F). Similar slow recovery was previously reported for IFA-1 and IFB-1 in the pharynx (Karabinos et al., 2003). In contrast, BBLN-1 recovered roughly 90% within the same period, indicating that BBLN-1 localizes to the IF network in a dynamic fashion. Taken together, these results indicate that BBLN-1 is a dynamic component of the endo-tube in intestinal cells.

### BBLN-1 requires an IF network for its localization and interacts with IF proteins

As BBLN-1 localizes together with IF proteins, we next determined whether the apical localization of BBLN-1 depends on IFs. The *C. elegans* intestine expresses six cytoplasmic IF proteins, all of which localize almost exclusively to the endotube (Bossinger et al., 2004; Geisler et al., 2020; Karabinos et al., 2002, 2004). We used RNAi feeding to knock down the individual intestinal IF proteins in a strain that endogenously expresses GFP::BBLN-1 and ERM-1::mCherry. We observed significant reduction in apical BBLN-1 upon knockdown of IFB-2, IFD-1, IFD-2, or IFP-1 (Fig. 3A, B). In *ifb-2*(*RNAi*) animals, the loss of BBLN-1 from the apical cortex was most striking. IFB-2 is an essential component of the apical intestinal IF network, and loss of IFB-2 results in a complete absence of the endotube (Geisler et al., 2019, 2020). Therefore, the strong reduction in apical BBLN-1 may be due to an additive effect of losing the localization of all cytoplasmic IFs. These data demonstrate that BBLN-1 requires the presence of an IF network for its apical localization. We also examined the localization of BBLN-1 in the excretory canal of *ifc-2*(*RNAi*) animals, as IFC-2 has been shown to be important for the structure of this tissue (Al-Hashimi et al., 2018). We observed a marked relocalization of BBLN-1 from the apical cortex to the cytoplasm and basal membrane, which suggests that dependency on IFs is a general aspect of BBLN-1 localization (Fig. 3C).

**Figure 3.**
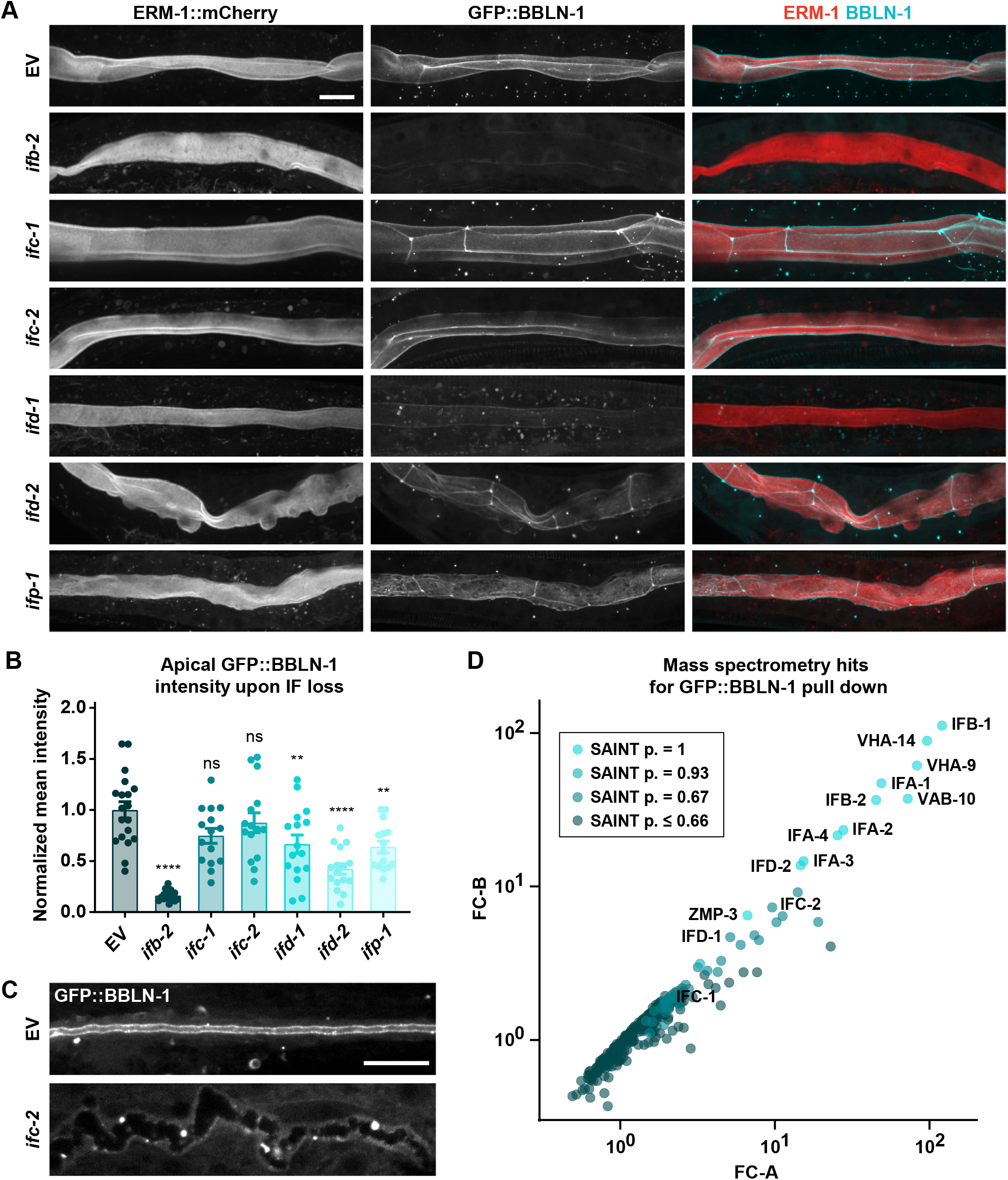
BBLN-1 requires intermediate filaments for its apical localization in the intestine. (**A**) Intestinal GFP::BBLN-1 distribution and apical membrane morphology, visualized by ERM-1::mCherry, upon RNAi knockdown of targets indicated to the left. EV = empty vector. (**B**) Quantification of GFP::BBLN-1 levels at the intestinal apical domain of larvae subjected to RNAi knockdown of targets indicated at the X-axis. Each data point represents the average of six or eight measurements in a single animal (n = 19, 17, 15, 15, 16, 16, and 15 animals, respectively). Larvae were L2–L4 stage. Data is presented as mean ± SEM and analyzed with ordinary one-way ANOVA followed by Dunnett’s multiple comparisons test; ns = P > 0.05, ** = P < 0.005, **** = P < 0.0001. (**C**) GFP::BBLN-1 distribution in the excretory canal upon RNAi knockdown of targets indicated on the left. EV = empty vector. (**D**) Mass spectrometry hits for GFP::BBLN-1 pull down plotted as correlation between fold-change (FC) score A and more stringent FC score B. Data points are color coded for different SAINT probability scores.

The reliance on an intact IF network for BBLN-1 to properly localize might originate from a physical interaction with IF proteins. To identify candidate interacting proteins, we performed affinity purification of GFP::BBLN-1 from mixed-stage *C. elegans* cultures, followed by mass spectrometry analysis. The highest-ranking proteins we identified included all intestinal IF proteins except IFP-1, as well as all proteins of the IFA/ IFB system (IFA-1, 2, 3, 4 and IFB-1) (Fig. 3D). After IFB-1, the two most highly ranked candidate interactors were VHA-9 and VHA-14, the *C. elegans* orthologs of subunits F and D of the stalk region of the highly conserved vacuolar ATPase (V-ATPase) V1 domain, which localizes to the luminal domain of the intestine (Ji et al., 2006). Other high-confidence hits included various hemidesmosome components (i.e. VAB-10, LET-805, PAT-12 and MUP-4) and actin-interacting proteins like the crosslinker FLN-2, and the myosins UNC-15 and MYO-2. Altogether, our mass spectrometry data suggests a physical interaction between BBLN-1 and IFs and provides cues for future investigations into the activities of BBLN-1.

To further confirm the dependency of BBLN-1 localization on IFs, we examined the effects of inactivation of known regulators of intermediate filament organization on BBLN-1. In addition to *sma-5*, loss of *ifo-1, act-5*, and *let-413* have all been reported to disrupt IFB-2 distribution in the *C. elegans* intestine (Bossinger et al., 2004; Carberry et al., 2012; Estes et al., 2011; Stutz et al., 2015). IFO-1 localizes to the adluminal domain, together with but independently from IFB-2, and its depletion causes aggregation of IFs at cell junctions as well as in the cytoplasm (Carberry et al., 2012). Reducing the levels of the intestinal actin ACT-5 causes gaps in the normally uninterrupted IFB-2 network, and has been reported to result in the appearance of cytoplasmic invaginations (Estes et al., 2011; Stutz et al., 2015). Finally, depletion of LET-413 causes aberrant localization of IFB-2 to the lateral and basal domains of intestinal cells in the embryo (Bossinger et al., 2004). We depleted each protein by RNAi in a strain expressing either ERM-1::mCherry (to visualize lumen morphology) or IFB-2::mCherry, together with GFP::BBLN-1. The phenotypes we observed matched published reports, though RNAi for *act-5* caused more severe IFB-2 defects, with invaginations only visible in a small subset of animals (Fig. 4A–C). Importantly, in each case the localization of BBLN-1 followed the localization pattern of IFB-2, confirming that BBLN-1 colocalizes with IFs.

**Figure 4.**
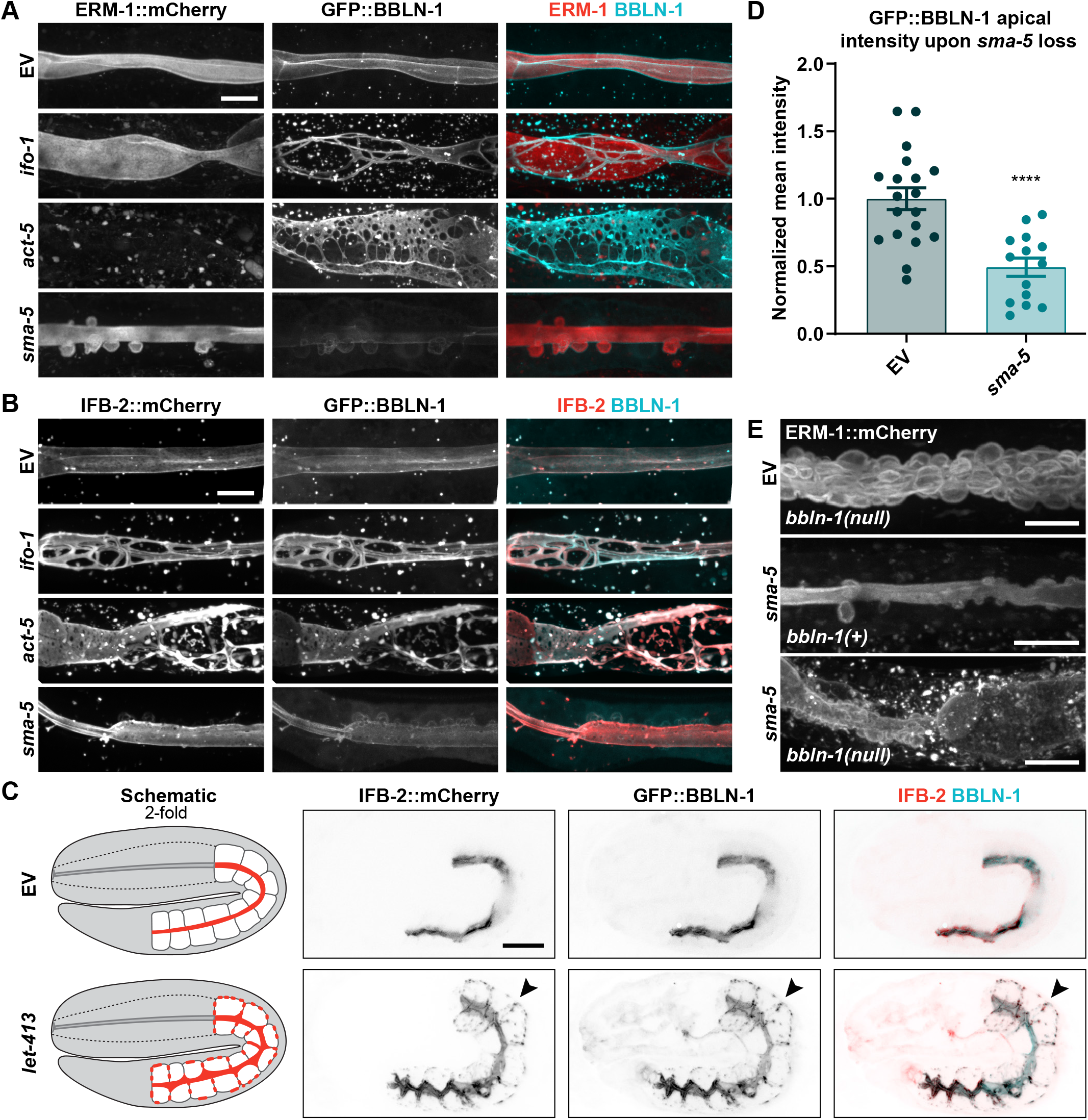
BBLN-1 co-localizes with disrupted or mislocalized IFs. (**A**) Intestinal GFP::BBLN-1 distribution and apical membrane morphology, visualized by ERM-1::mCherry, after RNAi knockdown of targets indicated on the left. EV = empty vector control. Same EV control is shown as in Fig. 3A since data is from a single experiment. (**B, C**) GFP::BBLN-1 and IFB-2::mCherry distribution in L2 (*act-5*), L3 (*ifo-1, sma-5*) and L4 (EV) larvae (B) and 2-fold embryos (C) after RNAi knockdown of targets indicated on the left. (**D**) Quantification of GFP::BBLN-1 levels at the intestinal apical domain of L3 and L4 larvae subjected to RNAi knockdown of *sma-5*. Each data point represents the average of six or eight measurements in a single animal (n = 19 and 14 animals, respectively). Data is presented as mean ± SEM and analyzed with ordinary one-way ANOVA followed by Dunnett’s multiple comparisons test; **** = P < 0.0001. EV control data is same as in Fig. 3B. (**E**) Apical membrane morphology, visualized by ERM-1::mCherry, after RNAi knockdown of targets indicated on the left in *bbln-1*(*+*) or *bbln-1*(*null*) background.

Interestingly, in *sma-5*(*RNAi*) intestines, we observed a significant reduction in cortical enrichment of BBLN-1 (Fig. 4D). This suggests that *sma-5* acts upstream of *bbln-1*. To determine if *sma-5* and *bbln-1* act in a linear pathway, we performed *sma-5* RNAi in a *bbln-1*(*null*) background. The resulting animals displayed more severe intestinal abnormalities than either *sma-5*(*RNAi*) or *bbln-1*(*null*) alone (Fig. 4E). Thus, while *sma-5* may act in part through *bbln-1*, both genes act at least partially independent. Collectively, our results indicate that the subcellular distribution of BBLN-1 depends physically on the presence of intermediate filaments.

### Loss of bbln-1 compromises the integrity of the IF network

Apical domain invaginations in *sma-5* mutants were previously linked to defects in IF organization by fluorescence and electron microscopy (Geisler et al., 2016). Unlike the uniform electron-dense endotube surrounding the lumen in wild-type animals, endotube thickness in *sma-5* mutant animals is highly variable (Geisler et al., 2016). The phenotypic similarities between *sma-5* and *bbln-1* mutants, as well as the localization of BBLN-1 to IF-rich structures, led us to hypothesize that apical domain invaginations in *bbln-1* mutant animals were similarly associated with defects in intestinal IF organization. To address this, we analyzed the distribution of three different intestinal IF proteins in *bbln-1*(*null*) animals: IFB-2, IFC-2, and IFD-2. For IFC-2, we used an existing CRISPR knock-in line expressing IFC-2a/e fused to the yellow fluorescent protein (IFC-2a/e::YFP) (Geisler et al., 2019, 2020). For IFD-2 we generated an endogenous *gfp::ifd-2* locus using CRISPR/Cas9.

In control animals, IFB-2 and IFC-2 were distributed evenly along the adluminal domain (Fig. 5A, C). Both proteins were also enriched at cell junctions, with IFC-2 showing stronger junctional enrichment than IFB-2. These localization patterns match prior observations (Hüsken et al., 2008; Carberry et al., 2012; Geisler et al., 2016, 2020). In contrast to IFB-2 and IFC-2, IFD-2 showed a clearly punctate localization pattern and was excluded from cell junctions (Fig. 5E, S2A). The distinct localization of IFD-2 indicates functional specialization within the IF network.

**Figure 5.**
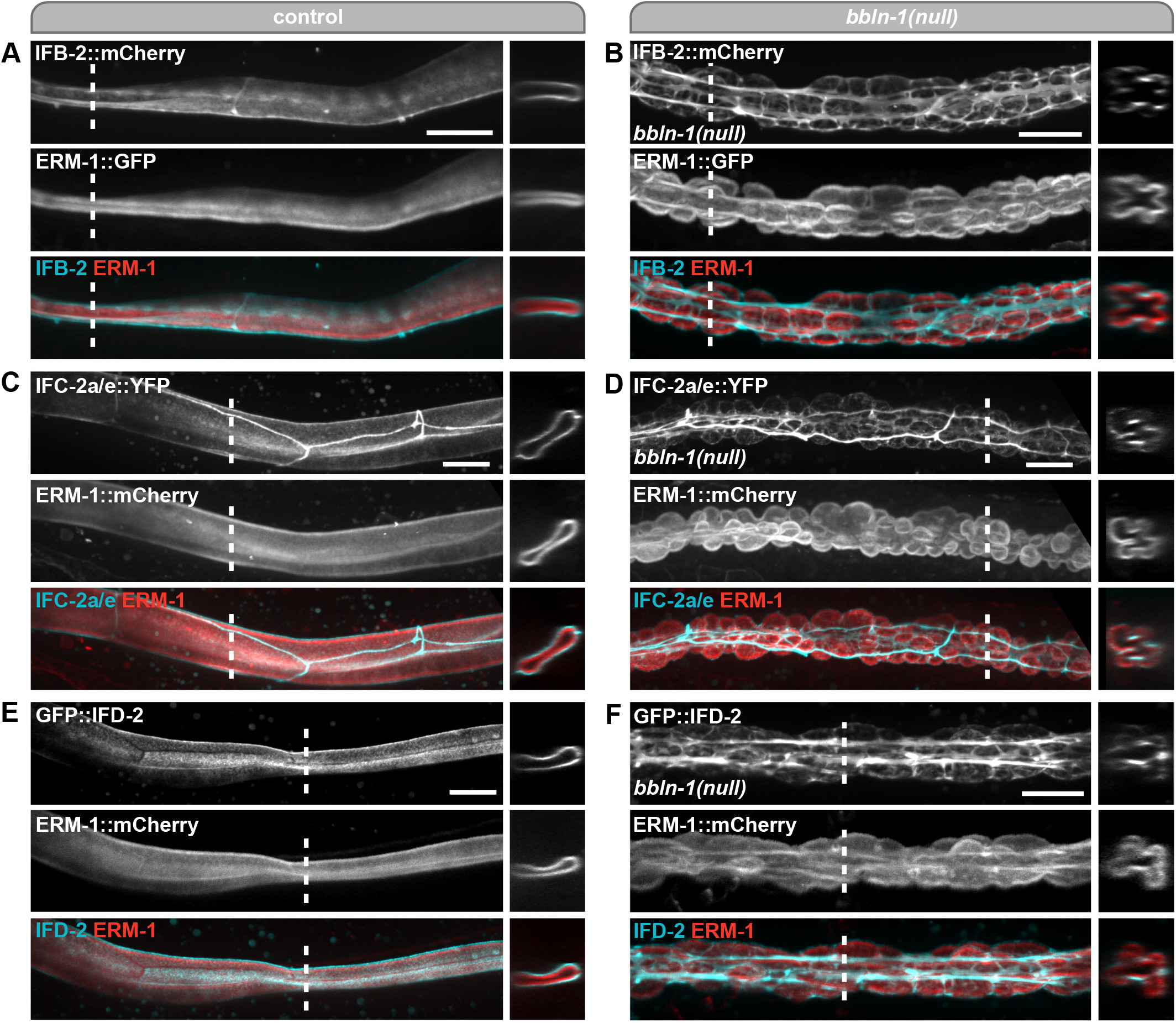
Loss of *bbln-1* compromises the integrity of the intermediate filament network. (**A–F**) Organization of the intestinal IF network visualized with IFB-2::mCherry (A, B), IFC-2a/e::YFP (C, D) or GFP::IFD-2 (E, F) in control (A, C, E) or *bbln-1*(*null*) (B, D, F) animals. Large panels are lateral views and small panels are cross-sections at the site indicated by the dashed lines.

Upon loss of *bbln-1*, all three IF proteins retained their subapical localization, but formed a network of cables or bundles of increased fluorescence intensity, surrounding protrusions into the cytoplasm with reduced fluorescence signal (Fig. 5B, D, F). IFD-2 lost its punctate localization pattern, instead localizing to cable-like structures (Fig. 5F). To investigate the stability of IFs within the aggregated bundles, we performed FRAP on IFB-2::mCherry in *bbln-1*(*null*) animals. IFB-2 showed very little fluorescence recovery, appearing if anything even more stable than in control animals (Fig. 2E, F). Together these data indicate that BBLN-1 is required for the integrity of the IF network but not for IF recruitment to the subapical cortex.

To investigate whether BBLN-1 is required for IF network integrity throughout development or only during a critical time in intestinal development, we made use of the auxin-inducible degradation (AID) system.The AID system enables targeted degradation of AID-degron tagged proteins through expression of the plant-derived auxin-dependent E3 ubiquitin ligase specificity factor TIR1 (Nishimura et al., 2009; Zhang et al., 2015). We used CRISPR/Cas9 genome engineering to tag BBLN-1 N-terminally with GFP and the AID-degron and expressed TIR1 from the intestine-specific *elt-2* promoter. Within 1 hour of addition of auxin to L1-stage animals, GFP::AID::BBLN-1 was no longer detectable at the luminal domain of the intestine (Fig. S2B). We then crossed the GFP::AID::BBLN-1 line with a line expressing IFB-2::mCherry and ERM-1::GFP to examine intestinal morphology over time. We added auxin to late L1 stage or early L4 stage animals and imaged the intestinal lumen after 24 h and 48 h (Fig. S2C). Irrespective of the starting time point, approximately half of the animals had developed invaginations after 24 hours of auxin treatment (Fig. S2D, 48-hour developmental timepoint). The remainder showed small ‘holes’ in the normally contiguous IFB-2::mCherry fluorescent pattern (Fig. S2E). Eventually, all animals treated with auxin from L1 developed cytoplasmic invaginations (Fig. S2D, 72-hour developmental timepoint). These data show that *bbln-1* is required to maintain the integrity of the IF network throughout development and further indicate that an intact IF network is essential for homeostasis of the intestinal lumen.

In addition to the IF network, the apical domain of the intestine is highly enriched in actin, which is present both in microvilli and the supporting terminal web. To investigate if the actin cytoskeleton is disrupted, we examined the localization of the intestinal actin ACT-5 in *bbln-1*(*null*) animals, using a YFP::ACT-5 transgene (Bossinger et al., 2004). In *bbln-1* animals, ACT-5 smoothly decorated the surface of the cytoplasmic invaginations and was not enriched in bundles or cables (Fig. S2F,G). Thus, loss of BBLN-1 function appears to specifically affect the IF network and not the terminal web or microvillar actin. We also investigated whether the loss of *bbln-1* disrupted the overall polarization of the intestinal cells. Inactivation of *bbln-1* by RNAi did not affect the basolateral localization of an endogenous LET-413/Scribble::mCherry marker, nor the apical localization of an endogenous PAR-6::GFP marker (Fig. S2H,I). Thus, the effects of *bbln-1* on the IF network are not due to a loss of intestinal polarity. Moreover, as invaginations are coated with PAR-6, ERM-1, and ACT-5, the formation of invaginations is not accompanied by a loss of apical identity.

To confirm loss of IF network integrity in *bbln-1* mutants and analyze intestinal morphology at the ultra-structural level, we performed electron microscopy on *bbln-1*(*null*) mutants. Previously, it was shown that in *sma-5* mutant intestines, the endotube is lost or diminished at sites of invagination and thickened around invagination necks (Geisler et al., 2016). In *bbln-1*(*null*) animals, the endotube appears to be similarly affected as in *sma-5* mutants. Endotube remnants appeared as aggregated electron-dense material, and no longer displayed the characteristic two lines of electron-dense material defining the endotube in wildtype intestines (Fig. 6, compare A vs. B or C vs. D). Nevertheless, as in wild-type animals, aggregated endotube material in *bbln-1*(*null*) intestines still maintained a characteristic distance from the apical membrane, consistent with the continued presence of the subapical terminal web (Fig. 6A, B, inlays). Finally, the electron microscopy images revealed that microvilli are still abundant in invaginations. These data suggests that the presence of BBLN-1 and a fully assembled endotube are not required for microvilli formation, which is consistent with previous data on *ifb-2* null mutants (Geisler et al., 2019).

**Figure 6.**
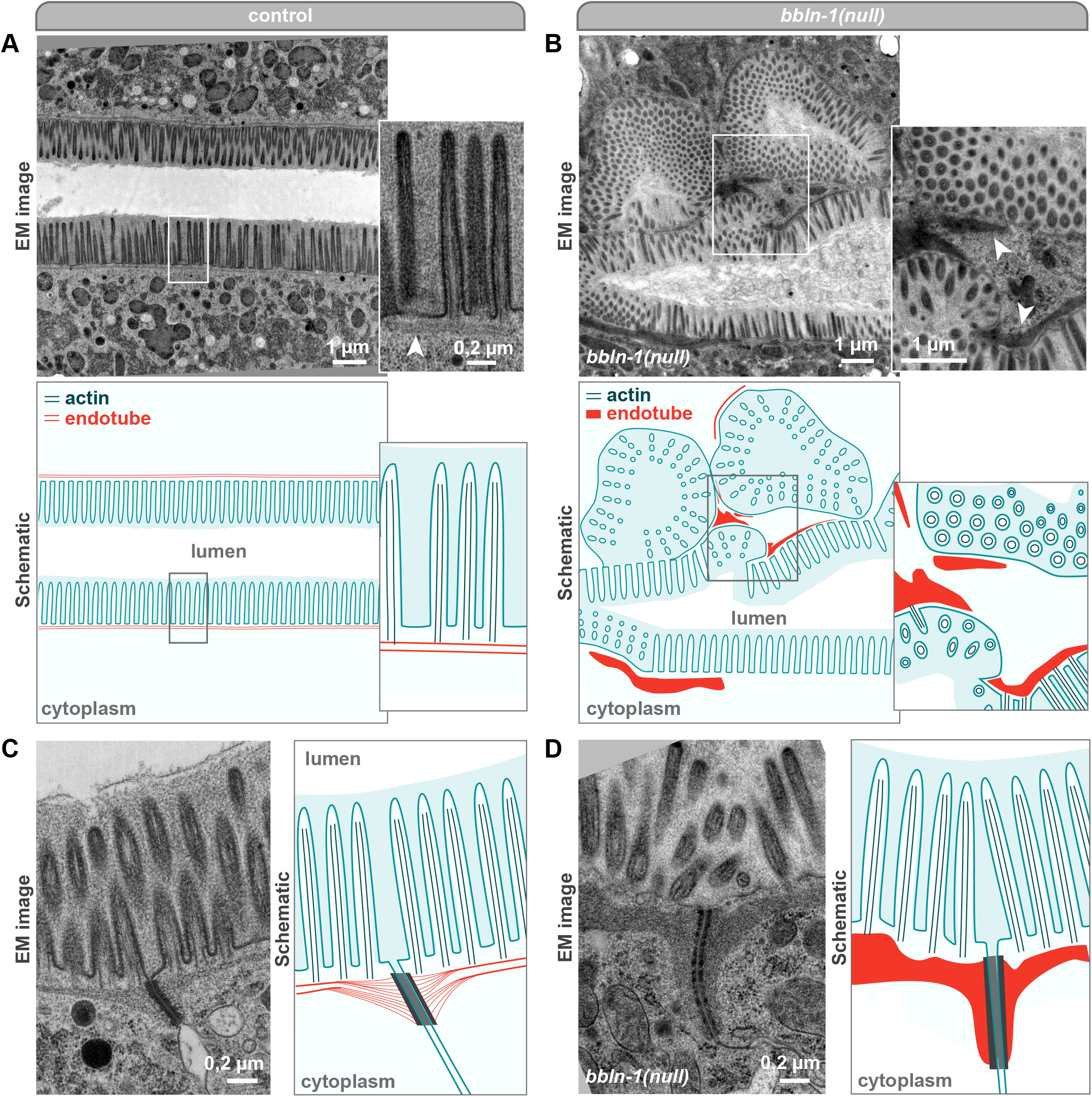
Loss of *bbln-1* compromises the integrity of the endotube. (**A, B**) Ultrastructure of the apical domain in intestinal cells of *bbln-1*(*+*) and *bbln-1*(*null*) adult animals visualized by transmission electron microscopy. Boxed region is shown enlarged and arrowheads point to the endotube. Schematics indicate actin bundles in blue and endotube in red. (**C, D**) Electron microscopy images showing the endotube at apical junctions. Schematics indicate actin bundles in blue, endotube in red, and junction in grey.

Taken together, the EM and light microscopy observations demonstrate that the intestinal IF network collapses and aggregates into stable bundles upon loss of BBLN-1, without affecting other aspects of intestinal epithelial polarity or the actin cytoskeleton.

### Apical IF aggregation drives cytoplasmic invaginations

We next sought to better understand the relationship between the cytoplasmic invaginations and collapsed IF network observed in *bbln-1* mutants. If invaginations are a consequence of the IF network collapse, more direct disruption of the IF network might be expected to similarly cause invaginations. Indeed, *ifd-2*(*RNAi*) induced cytoplasmic invaginations, though with much lower penetrance and severity than *bbln-1* mutants (Fig. 3A). A recent study similarly showed that *ifd-2* knockout causes cytoplasmic invaginations, while retaining apical IFB-2 and IFC-2 localization (Geisler et al., 2020). We also observed that knockdown of *ifp-1* causes an irregular apical surface and mildly fragmented BBLN-1 distribution (Fig. 3A). These data indicate that the cytoplasmic invaginations observed in *bbln-1* mutants are caused by defects in the IF network.

Surprisingly, despite the loss of the complete intestinal IF network, *ifb-2* knockout animals have been reported to be overall healthy in appearance, with only mild intestinal lumen morphology defects and increased susceptibility to stresses (Geisler et al., 2020). We also observed an irregular lumen morphology in only a subset of *ifb-2*(*RNAi*) animals (Fig. 3A, S3A). Importantly, loss of *ifb-2* does not result in cytoplasmic invaginations (Fig. 3A, S3A). This indicates that the invaginations observed in *bbln-1* mutants are due to an altered IF network state. If this is the case, loss of the IF network should suppress the invaginations in *bbln-1* animals. To test this, we generated a double mutant carrying the *bbln-1*(*null*) allele and the *ifb-2*(*kc14*) deletion allele (Geisler et al., 2019), expressing ERM-1::mCherry to mark the membrane. Indeed, *ifb-2*(*kc14*) suppressed the intestinal defects of *bbln-1*(*null*) (Fig. S3B). The lumen morphology in the double knockout appeared identical to that seen for IFB-2 depletion alone in *ifb-2*(*RNAi*) animals (Fig. 3A) as well as previously published for *ifb-2*(*kc14*) (Geisler et al., 2020).

In agreement with the fluorescence microscopy data, we did not observe cytoplasmic invaginations by electron microscopy in *ifb-2*(*kc14*); *bbln-1*(*null*) double mutants (Fig. S3C). No endotube was visible, confirming the requirement for IFB-2 in endotube formation. Surprisingly, we did observe occasional small membrane protrusions directed towards the lumen (Fig. S3D), which were also visible by light microscopy in both *ifb-2*(*kc14*); *bbln-1*(*null*) (Fig. S3E) and *ifb-2*(*RNAi*) animals (Fig. S3Aiii), and are therefore likely to be the result of IFB-2 depletion alone. Microvillar actin bundles appeared to cluster together at the neck of these small protrusions, forming fan-like structures. One possibility is that, in the absence of the supporting role of the endotube, microvillar actin bundles become linked together, causing the formation of these small extrusions.

Taken together, our data are consistent with a model in which loss of *bbln-1* leads to an altered, pathogenic IF network state that results in cytoplasmic invaginations.

### Bublin is the mammalian ortholog of BBLN-1

In both *C. elegans* and mammals, intestinal IFs function in mechanical support and protection against stresses. While the IF protein families have diverged between *C. elegans* and mammals (Erber et al., 1998; Weber et al., 1989), it is likely that essential aspects of the regulation of IF networks are conserved. We therefore investigated if BBLN-1 is conserved in mammals. An iterative search using Jackhmmer (Potter et al., 2018) revealed the uncharacterized protein C9orf16 as the only candidate mammalian homolog (E-value 3.8e-16). The 83 aa human C9orf16 protein is predicted to consist of a coiled-coil domain, flanked by intrinsically disordered regions — similar to the predicted structure of BBLN-1 (Fig. 7A). The region between amino acids 27 to 102 in BBLN-1 is most similar to C9orf16, with 26% shared amino acid identity and 48% similarity. A reciprocal Jackhmmer search starting with C9orf16 also identified BBLN-1 as the only *C. elegans* homolog (E-value 8.4e-07). We used the DRSC Integrative Ortholog Prediction Tool v7.1 (Hu et al., 2011) to query multiple orthology prediction databases, but only OrthoDB v9 (Zdobnov et al., 2017) identified C9orf16 and BBLN-1 as candidate orthologs. Nevertheless, given the bi-directional best hit in Jackhmmer and the absence of other candidate orthologs, we consider it likely that BBLN-1 and C9orf16 are evolutionary orthologs and we have therefore named the human gene product bublin coiled-coil protein (BBLN).

**Figure 7.**
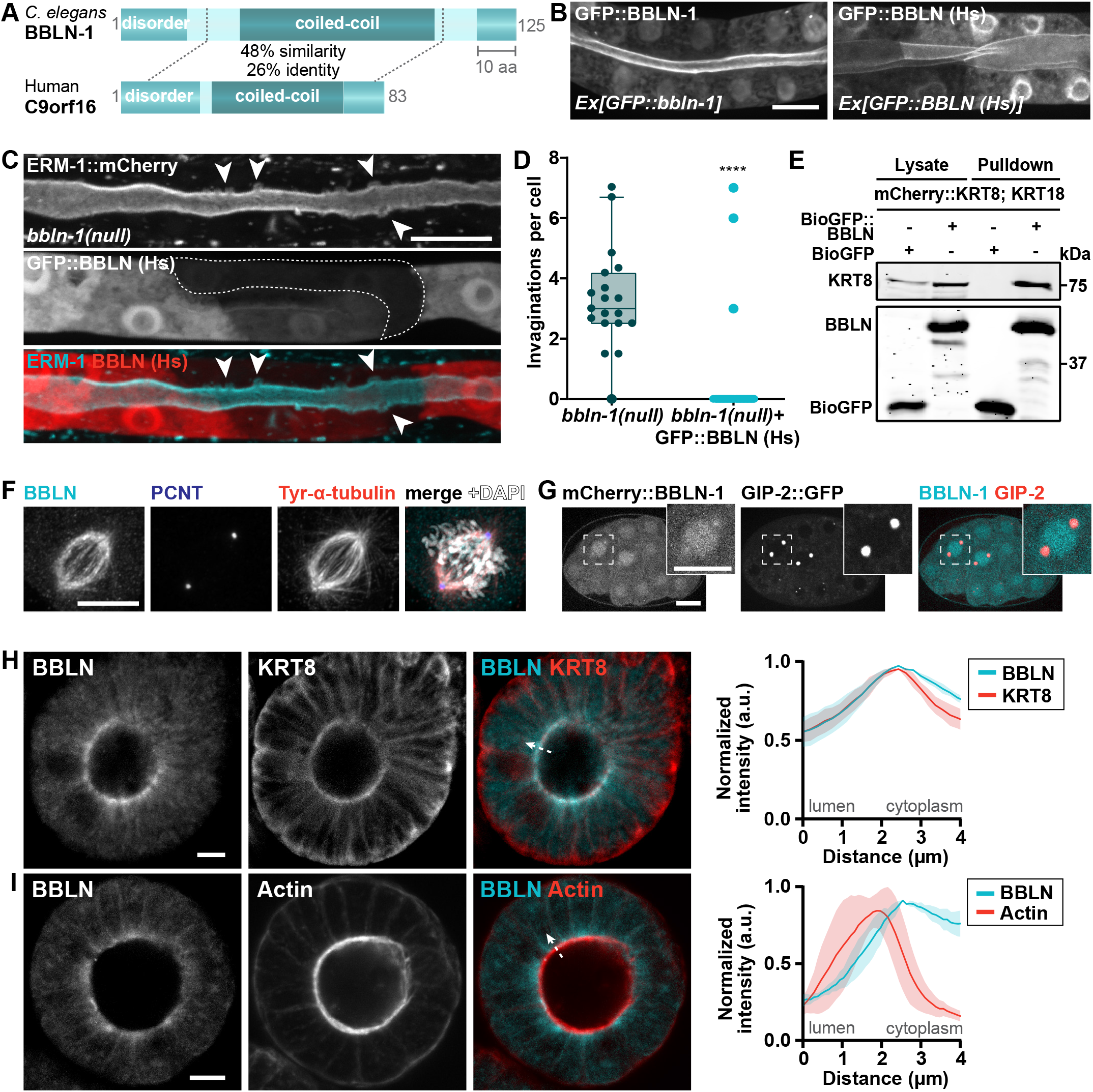
Bublin is the mammalian homologue of BBLN-1. (**A**) Schematic representation of *C. elegans* BBLN-1 and human C9orf16 proteins with predicted domains. EMBOSS Water analysis calculates 26% identity and 48% similarity over a length of 77 aa. (**B**) Overexpression of GFP-tagged BBLN-1 and bublin (BBLN (Hs)) in *C. elegans* under control of the intestine-specific *vha-6* promoter. (**C**) Apical membrane morphology visualized by ERM-1::mCherry in a *bbln-1*(*null*) L1 larva with mosaic expression of GFP tagged bublin (BBLN (Hs)) from an extrachromosomal array. The cell without GFP::BBLN (Hs) expression (dotted outline) displays cytoplasmic invaginations (arrowheads). (**D**) Quantification of the number of invaginations in intestinal cells in *bbln-1*(*null*) L1 larvae that either express or do not express intestinal GFP::BBLN (Hs). Each dot corresponds to a single cell (19 cells over 8 animals for control, 38 cells over 16 animals for the rescue). Data is presented as median ± interquartile range with Tukey whiskers, and data is analyzed with Mann-Whitney test; **** = P < 0.0001. (**E**) Western blots of Biotin pull-down from extracts of HEK293T cells transfected with BirA, mCherry::KRT8, KRT18 and either BioGFP::BBLN or BioGFP alone as control. Blots were probed for keratin 8 (KRT8, top) and GFP (bottom). The input is 10% of the biotin pull-down. (**F**) Point-scanning confocal microscopy images of HeLa cell stained with antibodies for bublin (BBLN), pericentrin (PCNT) and tyrosinated α-tubulin. (**G**) Endogenous mCherry::B-BLN-1 and GIP-2::GFP distribution during cell division in the early *C. elegans* embryo. (**H, I**) Section of mouse small intestinal organoid stained with antibodies against bublin and keratin 8 (KRT8, top) or with phalloidin to visualize actin (bottom). Images were acquired using point-scanning confocal microscopy. Graphs to the right show the fluorescence intensity from lumen to cytoplasm (example indicated with the dotted arrow). Fluorescent intensity was measured across all cells that are in focus, and displayed as mean ± SD.

To investigate if bublin and BBLN-1 are functionally conserved, we generated transgenic animals expressing GFP::bublin in the intestine of *C. elegans* from an extrachromosomal array. Bublin showed apical and perinuclear enrichment, resembling the localization pattern of BBLN-1 when similarly expressed from an extrachromosomal array (Fig. 7B). Moreover, as we observed for BBLN-1, the sub-apical localization of bublin was dependent on the presence of an intact IF network (Fig. S4A). To determine if bublin can functionally replace BBLN-1, we examined the effects of mosaic GFP::bublin expression in a *bbln-1*(*null*) background. As expected, *bbln-1*(*null*) animals not inheriting the transgenic array all showed invaginations from hatching (Fig. 7D). In contrast, in mutant animals that did inherit the transgene, cells expressing GFP::bublin did not show invaginations at hatching (Fig. 7C, D). As animals developed, invaginations began to form in all cells, and animals became phenotypically indistinguishable from non-transgenic animals by mid-L1 stage. Thus, expression of bublin delays the invagination phenotype in young larvae. Together, these findings indicate that bublin is a functional homolog of BBLN-1.

We next investigated if the interaction with intermediate filaments is conserved. Keratin 8 (KRT8) is the most common type II keratin IF in simple epithelia, and in enterocytes keratin 8 is part of an apical IF network (Schwarz et al., 2015). As keratin 8 forms functional heterodimers with keratin 18 (KRT18) (Asghar et al., 2015), we investigated if bublin interacts with keratin 8/18 by affinity purification from HEK293T cells. Pull-down analysis from HEK293T cell lysates showed that keratin 8/18 co-purify with bublin (Fig. 7E). Additionally, multiple high-throughput studies found bublin as an interactor of the type I intestinal keratin 20 (Luck et al., 2020; Rolland et al., 2014; Rual et al., 2005). Thus, the interaction with intermediate filaments is conserved between BBLN-1 and bublin.

We also performed mass spectrometry analysis on bublin purified from HEK293T cells. The top scoring hit was V-ATPase subunit D (ATP6V1D), the ortholog of *C. elegans* VHA-14 and second highest scoring hit in our BBLN-1 interaction screen (Fig. S4B, 4D). We additionally found subunit F of this complex (ATP6V1F), the ortholog of the high-confidence BBLN-1 interactor VHA-9. Thus, the similarity in interactions between BBLN-1 and bublin extends to V-ATPase subunits, further strengthening that bublin is the ortholog of BBLN-1.

Finally, we investigated the subcellular localization of bublin in mammalian cells using a GFP::bublin expression construct and a polyclonal antibody directed against human bublin. We first confirmed specificity of the antibody by immunostaining HeLa cells transfected with a GFP::bublin expression construct. The staining and GFP fluorescence patterns overlapped, demonstrating that the antibody recognizes bublin (Fig. S4C). We next stained HeLa cells with antibodies recognizing bublin and keratin 8. However, we did not detect overlap between the bublin and keratin 8 localization patterns (Fig. S4D). Thus, whereas *C. elegans* BBLN-1 appears to exclusively co-localize with intermediate filaments, this is not the case for bublin. Interestingly, in dividing HeLa cells, bublin localized to interpolar and kinetochore microtubules, a localization we had not observed for BBLN-1 (Fig. 7F). We therefore re-examined the localization of BBLN-1 in *C. elegans* embryonic mitotic cells also expressing the γ-tubulin interacting protein GIP-2::GFP to mark the centrosomes. BBLN-1 was enriched at the interpolar region (Fig. 7G), indicating that both BBLN-1 and bublin operate at the mitotic spindle.

HeLa cells do not contain the subapical IF web present in *C. elegans* intestinal cells or in differentiated enterocytes. We reasoned that the lack of co-localization between bublin and Keratin 8 might reflect the absence of this particular IF structure. We therefore examined the localization of bublin in mouse small intestinal organoids, which mimic the normal structure and composition of the intestinal epithelium much more closely than traditional cell cultures. In organoids, bublin showed clear enrichment at the apical domain of intestinal epithelial cells. Bublin localized just below the apical actin network, at the same level as Keratin 8 (Fig. 7H, I). Thus, the localization of bublin in mouse intestinal organoids mimics that of BBLN-1 in *C. elegans* intestinal cells. Collectively, the overlap in localization, rescue of phenotype, and similarity in interaction partners, suggest that bublin and BBLN-1 have at least partially overlapping functionality.

## Discussion

Formation and maintenance of tubular architecture is a complex and dynamic morphogenetic process that relies on apically localized structural components conserved across animal species. Using an unbiased approach in the animal model *C. elegans* we identified the small protein BBLN-1 as a regulator of tubular architecture of the intestinal epithelium. Loss of BBLN-1 causes bubble-shaped invaginations of the apical plasma membrane into the cytoplasm, characterized by aggregation of IFs into a network of fibers that outline these protrusions. Ultrastructurally, loss of BBLN-1 results in endotube thickening between invaginations and endotube loss at invaginating membrane regions. In contrast to the severe defects in IF organization, actin, cell polarity markers, and cell junctions are largely unaffected. Similar observations were made in a dissertation that describes the identification of a mutant carrying the same point mutation present in *bbln-1*(*mib42*) (Paulson, 2009).

We identified C9orf16 as the mammalian ortholog of BBLN-1 based on sequence homology and have named it bublin coiled-coil protein (BBLN). When expressed in the intestine of *C. elegans*, the localization pattern of bublin mimicked that of BBLN-1, including the IF-dependent apical localization. Moreover, expression of bublin was able to partially suppress the *bbln-1* phenotype, delaying the appearance of invaginations. Affinity purification and mass spectrometry experiments also showed a strong overlap in candidate interaction partners between BBLN-1 and bublin. Finally, in mouse intestinal organoids we observed localization of bublin to the apical IF network, just below the actin-rich terminal web. All of these observations are consistent with bublin being the ortholog of BBLN-1. Nevertheless, while BBLN-1 seems to strictly co-localize with IFs in *C. elegans*, this is not the case for bublin, as we did not observe co-localization with keratin 8 in HeLa cells. Thus, while bublin may regulate IF organization, its functioning may have diverged between *C. elegans* and mammals.

Multiple observations support that BBLN-1 is a direct regulator of IFs, and that the cytoplasmic invaginations in *bbln-1* mutants are a consequence of a disorganized IF network organization. First, BBLN-1 colocalizes with IFs, in a manner dependent on the presence of an IF network. Multiple IF proteins co-purified with BBLN-1 from *C. elegans* lysates, indicating that the localization of BBLN-1 is mediated through physical interactions. Second, phenotypes similar to *bbln-1*(*null*) have been described for mutants of the IF member *ifd-2*, and for *sma-5*, a regulator of the IF network whose loss causes cytoplasmic invaginations that are strikingly similar to those present in *bbln-1* mutants (Carberry et al., 2012; Geisler et al., 2016, 2020). Finally, and most strikingly, loss of the apical IF network through knockout of *ifb-2* suppressed the *bbln-1* phenotype, demonstrating that the presence of apical IFs is a prerequisite for invagination formation. The most likely explanation is that *bbln-1* causes an altered, pathogenic IF network state that leads to the formation of invaginations into the cytoplasm.

Why a compromised IF network leads to cytoplasmic invaginations is not clear. One explanation is that pressure from the lumen physically forces invaginations through weak spots in the IF network. However, *ifb-2* mutants which lack an apical IF network entirely, have a largely normal intestinal morphology and do not show lumen widening until the L4 stage (Geisler et al., 2020). Possibly the actin network compensates for the loss of the IF network and delimits luminal widening. The presence of compensatory mechanisms involving actin is supported by the observation that loss of *ifb-2* leads to an increase in apical levels of the actin-bundling protein plastin 1 (PLST-1) (Geisler et al., 2020). Another explanation for the formation of invaginations is that the IF network plays a more direct instructive role in determining the width of the lumen or the surface area of the apical membrane. Such a role has recently been proposed for the excretory canal, where IFs work in conjunction with actin and tubulin to control intracellular lumenogenesis (Khan et al., 2019). Here, IFs are hypothesized to regulate access of membrane formation promoting vesicles to the lumen, to restrain lateral lumen expansion. Defects in apical membrane trafficking in the intestine have been shown to cause ectopic lateral lumen formation (Shafaq-Zadah et al., 2012). Hence, it is possible that the IF network plays a role in apical trafficking in the intestine.

The molecular nature of the change to the IF network in *bbln-1* and the phenotypically very similar *sma-5* mutants also remains to be determined. SMA-5 is a stress-activated MAPK homolog, and its loss results in increased phosphorylation of IFs. Hence it likely exerts an indirect effect on IFs, but relevant downstream targets have not been identified. We therefore considered the possibility that BBLN-1 acts downstream of SMA-5. However, the increased phenotype severity we observed when *sma-5* was depleted in the *bbln-1*(*null*) background suggests that these genes do not act in a simple linear pathway. The protein sequence of BBLN-1 does not clearly hint at its function, as it lacks functionally distinct domains. Possible roles for BBLN-1 include cross-linking or stabilizing IF filaments, and anchoring of IFs to components of the overlying terminal web. The tendency of invaginations to occur at cell junctions may also indicate a role in linking IFs to cell junctions. However, our FRAP data indicates that BBLN-1 associates dynamically with the IF layer. Hence, the activities of BBLN-1 may be more indirect, for example localizing unidentified regulators to the IF network or promoting post-translational modifications of IF proteins. Our interaction studies identified subunits of V-ATPases as prominent candidate interactors of BBLN-1 and bublin. Whether these interactions are physiologically relevant and if they are involved in IF regulation or other aspects of BBLN-1 function remains to be determined. A final intriguing hypothesis is that BBLN-1 is involved in protein aggregation. Bublin was found in a search for highly heat-resistant proteins that remain soluble upon boiling (Tsuboyama et al., 2020). The identified proteins, termed ‘Hero’ for heat-resistant obscure proteins, were shown to protect different subsets of proteins from aggregation or denaturation under stress conditions (Tsuboyama et al., 2020). Indeed, bublin is differentially expressed in diseases characterized by protein aggregation, including amyotrophic lateral sclerosis (ALS) (Nijssen et al., 2018), Alzheimer’s (Kong et al., 2009), and Parkinson’s disease (Kim et al., 2020). As the loss of BBLN-1 leads to IF aggregation, a potential future avenue of investigation is to examine if BBLN-1 plays a direct role in protecting against protein aggregation.

In summary, we identified a conserved regulator of IF organization, whose functioning highlights the importance of IF network integrity in maintaining regular lumen shape. While our study has focused on the role of BBLN-1 in organization of the apical IF network of intestinal cells, it is likely that BBLN-1 plays additional roles in different tissues and processes. The BBLN-1 localization pattern in other tissues strongly resembled that of IFs, and the sub-apical localization of BBLN-1 in the excretory canals depended on IFC-2. Thus, BBLN-1 may be a broad regulator of IF organization in *C. elegans*.

## Supporting information

Supplemental DNA files

## Acknowledgments

We thank R. Schmidt for sharing strain BOX459, V. Garcia Castiglioni for strains BOX251 and BOX260, K. Oegema for strain OD2509, J. Sepers for the NRFL-1::mCherry fusion, and B. van der Vaart for the keratin plasmids. We thank M. Kersten, J.-P. ten Klooster, W. Nijenhuis, and N. Schwarz for technical assistance. We thank S. van den Heuvel, S. Ruijtenberg, M. Harterink, B. Mulder, B. Snel and members of the Van den Heuvel and Boxem groups for helpful discussions. We also thank WormBase, the HUGO Gene Nomenclature Committee (HGNC) and the Biology Imaging Center, Faculty of Sciences, Department of Biology, Utrecht University. Some strains were provided by the Caenorhabditis Genetics Center, which is funded by NIH Office of Research Infrastructure Programs (P40 OD010440).

## Funding

This work was supported by the Netherlands Organization for Scientific Research (NWO)-CW ECHO 711.014.005 and NWO-VICI 016.VICI.170.165 grants to M. Boxem, and the German Research Council (LE566/14-1, 3; R. Leube). This research was part of the Netherlands X-omics Initiative and partially funded by NWO, project 184.034.019.

## Author Contributions

Conceptualization: S.R., J.J.R., M.B.; Formal analysis: S.R., R.S., M.B.; Investigation: S.R., F.G., R.S., J.R.K., S.v.d.H., O.D.J., M.P., J.J.R.; Resources: S.v.d.H., A.A., C.A.R., M.A., R.E.L., M.B.; Data Curation: S.R., J.R.K.; Writing – Original Draft: S.R., J.J.R., M.B.; Writing – Review & Editing: F.G., A.A., C.A.R., R.E.L.; Visualization: S.R., M.B.; Supervision: S.R., F.G., A.A., C.A.R., M.A., R.E.L., J.J.R., M.B.; Project Administration: M.B.; Funding Acquisition: A.A., C.A.R., M.A., R.E.L., M.B.

## Declaration of Interests

The authors declare no competing interests.

## Materials and Methods

### Caenorhabditis elegans strains and culture conditions

*Caenorhabditis elegans* strains were cultured under standard conditions (Brenner, 1974). All experiments were performed with animals grown on nematode growth medium (NGM) agar plates at 20 °C. Table S1 contains a list of all the strains used. Transgenic lines were generated by injecting 15 ng/µl of either plasmid pSMR10 or pSMR30 together with 65 ng/µl lambda DNA (Thermo Scientific) into the gonads of young adults.

### Cell line culture

HeLa and HEK293T cell lines were cultured in DMEM/Ham’s F10 (50:50) supplemented with 10% FCS and 1% penicillin/streptomycin at 37 °C and 5% CO2. All cell lines routinely tested negative for mycoplasma.

### Organoid culture

Mouse small intestinal organoids derived from the duodenum of C57BL/6 mice were cultured in ENR medium: Advanced DMEM/F12 (Invitrogen) with 1% Penicillin/Streptomycin (P/S, Lonza), 1% Hepes buffer (Invitrogen) and 1% Glutamax (Invitrogen), supplemented with 5% R-spondin conditioned medium, 10% Noggin conditioned medium, 50 ng/ml EGF (Invitrogen), 1x B27 (Invitrogen), 1.25 mM n-Acetyl Cysteine (Sigma-Aldrich).

### Isolation and initial mapping of mib41 and mib42

L4 stage *erm-1::GFP* animals were incubated for 6 hours at room temperature (RT) in M9 buffer (0.22 M KH_2_PO_4_, 0.42 M Na_2_HPO_4_, 0.85 M NaCl, 0.001 M MgSO_4_) supplemented with 50 mM ethyl methanesulfonate (EMS), in a 15 ml tube with gentle rotation. Mutagenized L4 animals were placed on 9 cm NGM agar plates seeded with *E. coli* (35 plates, 10 animals per plate) and allowed to lay eggs. Parents were removed after ∼1000 F1 progeny had been produced. Three days after F1 progeny started egg-laying, all F1 adults and L2 larvae were washed off the plate, leaving behind a semi-synchronous population of F2 embryos. F2 animals were allowed to develop for 2–4 days at 20 °C and scored for intestinal morphology defects using a Leica MZ16 fluorescence stereoscope. Two mutants, *mib41* and *mib42*, were isolated and backcrossed with N2 males 3 times before further analysis. To identify an approximate genetic location for both mutations, single-nucleotide polymorphism (SNP) mapping using the polymorphic strain CB4856 was performed as described previously (Davis et al., 2005).

### Genomic DNA purification

*C. elegans* lysis and genomic DNA purification were performed using the DNeasy Blood and Tissue kit (Qiagen). Animals from two recently starved populations were collected, pooled, washed twice using M9, and resuspended in 400 μl of lysis (ATL) buffer. Samples were then flash-frozen in liquid nitrogen and stored overnight at −80 °C. After three rounds of liquid nitrogen freeze-thawing, proteinase K was added to a final concentration of 2 mg/ml and samples were incubated for 3 hours at 56 °C while shaking at 600 rpm. Samples were further incubated with 2 mg/ml RNAse A for 30 min at RT and for 10 min at 56 °C after addition of 400 μl of AL buffer. Finally, samples were mixed with 400 μl of 100% ethanol and loaded into purification columns. Wash and elution steps were performed according to the manufacturer’s instructions. Concentration of genomic DNA samples was measured using Qubit Fluorometric Quantification (Invitrogen).

### Identification of mib41 and mib42 mutations

We used the sibling subtraction method to identify the causative *mib41* and *mib42* mutations by whole-genome sequencing (WGS) (Joseph et al., 2018). From heterozygous *erm-1::GFP/+; mib41/+* or *erm-1::GFP/+; mib42/+* hermaphrodites we isolated homozygous *mib41* or *mib42* mutant, homozygous nonmutant, or heterozygous F2 progeny. Genomic DNA was then purified from pooled F3 progeny derived from 12–16 F2 animals and sequenced on an Illumina HiSeq X Ten platform. Candidate causative variants were defined as present at a frequency >0.8 in the homozygous mutant sample, <0.8 in the heterozygous sample, and <0.1 in the homozygous wild-type sample. For both mutant alleles, only a single variant from the resulting selection was predicted to affect a protein coding sequence, which was selected for further analysis. See Supplemental methods for a complete description of the bioinformatics pipeline to identify variants.

### Protein structure and domain predictions

General protein domain searches were done using the SMART service at http://smart.embl-heidelberg.de/. Coiled-coils predictions were done using DeepCoil at https://toolkit.tuebingen.mpg.de/#/tools/deepcoil. Disorder predictions were done using DisEMBL at http://dis.embl.de/. Default settings were used.

### Phylogenetic analysis

To identify proteins related to C15C7.5/BBLN-1, we performed a three-iteration HMMER search against the Reference Proteomes dataset at http://www.ebi.ac.uk/Tools/hmmer/.

### Molecular cloning

SapTrap assembly was done as described (Schwartz and Jorgensen, 2016) using existing SEC donor modules (Dickinson et al., 2018) or new donor modules generated by cloning PCR fragments or custom gBlocks (IDT) into Eco53kI-digested vector pHSG298 (Takara Biosciences). For the *Pvha-6::GFP::bbln-1::tbb-2 3’UTR* and *Pvha-6::GFP::BBLN* (*Hs*)*::tbb-2 3’UTR* rescue constructs, *Pvha-6* was amplified from genomic N2 DNA, *GFP* and the *tbb-2 3’UTR* were amplified from prior plasmids, and the *bbln-1* and *bublin* sequences were synthesized as gBlocks (IDT). GFP sequence is codon-optimized and contains 3 artificial introns. A list of all used oligonucleotides (IDT) and gBlocks (IDT) is included in Table S2 and used plasmids are listed in Table S3. PCR fragments were generated using Q5 Hot Start High-Fidelity DNA Polymerase (New England Biolabs) and gel purified using the Nucleospin kit (Machery-Nagel). DNA concentrations were measured using a BioPhotometer D30 (Eppendorf). All DNA vectors used for genome editing were purified from DH5α *E. coli* using a Qiagen midiprep kit. Annotated DNA files of all plasmids used are included as Supplemental DNA files.

### CRISPR/Cas9 genome engineering

Endogenous gene fusions were generated in an N2 background by homology-directed repair of CRISPR/ Cas9-induced DNA double-strand breaks (DSBs). Microinjection of young adult hermaphrodite germlines was done using an inverted microinjection setup (Eppendorf FemtoJet 4x mounted on a Zeiss Axio Observer A.1 equipped with an Eppendorf Transferman 4r). In cases where the sgRNA target site was not disrupted by sequence integration, silent mutations were incorporated to prevent repeated DNA cleavage. In all cases correct integration was confirmed by Sanger sequencing (Macrogen). A list of all DNA- and RNA-based reagents is included as Table S2.

*erm-1::mCherry* was generated using the SEC method, using a plasmid-based repair template. sgRNA expression plasmids were generated by ligating annealed oligo pairs into BbsI-digested pJJR50 as previously described (Waaijers et al., 2016). The repair template was assembled into pMLS257 (Addgene #73716) using SapTrap with custom and SEC modules as follows (from 5’ to 3’): left homology arm, a C-terminal linker (pMLS287), mCherry, SEC (pDD363), auxin-inducible degron (AID; pDD398), and right homology arm. Homology arms of ∼600 bp upstream and downstream of the DSB were amplified from N2 genomic DNA. The injection mix contained: 60 ng/µl *Peft-3::Cas9* (Addgene #46168), 15 ng/µl repair template, 100 ng/ µl for each sgRNA, and 2.5 ng/µl *Pmyo-2::mCherry* (Addgene #19327). Three injected animals were pooled and incubated for 3 days at 20 °C before adding 250 ng/µl of hygromycin per plate. Rol animals lacking visible mCherry expression were selected after 4–5 days. To eliminate the selection cassette through Cre-Lox recombination, L1 progeny of selected homozygous Rol animals were heat-shocked in a water bath at 34 °C for 1 hour. Correct excision was confirmed by selection of non-Rol animals and subsequent Sanger sequencing. Sequence files of the final gene fusions are included in Supplemental DNA files.

The *bbln-1*(*mib70*), *GFP::bbln-1*, and *ifb-2::mCherry* alleles were generated using the plasmid-free nested-CRISPR approach (Vicencio et al., 2019). Injection mixes contained the following reagents (IDT): 250 ng/ µl Alt-R S.p. Cas9 Nuclease V3 (IDT), 2 µM step 1 ssODN repair template, 400 ng/µl step 2 PCR repair template, 4.5-5 µM step 1 and step 2 crRNAs, 10 µM tracrRNA, as well as 1 µM *dpy-10* crRNA and ssODN repair for co-CRISPR selection (Arribere et al., 2014). Reagents for *dpy-10* co-CRISPR selection were omitted in *ifb-2::mCherry* injection mixes, and for *bbln-1*(*mib70*) only step 1 editing events were selected. To select for integration events, injected animals were transferred to individual plates and allowed to recover at 20 °C overnight before incubation at 25 °C for 2–3 days. F1 animals were either visually screened for presence of fluorescence using a Leica MZ16 fluorescence stereomicroscope, or singled from plates with high numbers of Dpy and Rol animals followed by PCR screening.

All other strains were produced using a plasmid-free protocol incorporating melting of double stranded DNA (dsDNA) repair templates (Ghanta and Mello, 2020). Selection of positive editing events was done by visually screening for expression of fluorescent protein, or by PCR analysis of single animals from plates with high numbers of Rol animals.

### Light microscopy

Imaging of *C. elegans* was done by mounting embryos or larvae on a 5% agarose pad in a 10 mM Tetramisole solution in M9 buffer to induce paralysis. Spinning disk confocal imaging was performed using a Nikon Ti-U microscope equipped with a Yokogawa CSU-X1-M1 confocal head and an Andor iXon DU-885 camera, using a 60x 1.4 NA objective. Time-lapse imaging for FRAP experiments was performed on a Nikon Eclipse-Ti microscope equipped with a Yokogawa CSU-X1-A1 spinning disk and a Photometrics Evolve 512 EMCCD camera, using a 100x 1.4 NA objective. Targeted photobleaching was done using an ILas system (Roper Scientific France/ PICT-IBiSA, Institut Curie). Point-scanning confocal microscopy was done on a Zeiss Airy-Scan LSM 880 setup using a Plan-Apochromat 63x 1.2 NA objective. bublin antibody test was imaged on an upright fluorescence Nikon Eclipse Ni-U microscope using a Plan Apo Lambda 100x N.A. 1.45 oil objective. Confocal imaging for keratin 8 and bublin in HeLa cells was performed with Leica TCS SP8 STED 3X microscope using HC PL APO 100x/1.4 oil STED WHITE objective driven by LAS X controlling software. Microscopy data was acquired using MetaMorph Microscopy Automation & Image Analysis Software (Spinning Disk), and Zen Black software (AiryScan). All stacks along the z-axis were obtained at 0.25 μm intervals. For quantifications, the same laser power and exposure times were used within experiments.

### Image analysis

Invagination numbers in *bbln-1*(*null*) animals were counted manually from intestinal rings int2 to int4, using Z-stacks that span the entire intestinal lumen. Invagination widths were measured by determining the full width at half-maximum of an intensity plot drawn across the center of the invagination in an orthogonal view. Corresponding cell lengths were quantified by drawing a spline through the lumen from lateral membrane to lateral membrane. Intensity distribution profiles of fluorescent proteins at the apical domains of the intestine or organoids were obtained by performing a 40-pixel (4.7 µm) wide line-scan perpendicular to the apical membrane. GFP::BBLN-1 intensity at the apical domain was determined by taking the maximum intensity value of a 40-pixel wide line-scan perpendicular to the apical membrane, and subtracting the mean background intensity (measured in a 40-pixel diameter circle outside of the worm). Each data point shown represents the average of 6–8 measurements per animal. All images were analyzed and processed using ImageJ alone (spinning disk) or in combination with Zen Black software (Airyscan).

### Quantification of brood size and lethality

Starting at the L4 stage, individual P0 animals were cultured at 20 °C and transferred to a fresh plate every 24 hours for 6 days. Hatched and unhatched progeny were scored 24 hours after removal of the P0, and larval lethality was scored 48 hours after removal of the P0.

### Texas Red-dextran assay

Mixed-stage populations were collected in egg buffer (118 mM NaCl, 48 mM KCl, 2 mM MgCl_2_, 2 mM CaCl_2_, 25 mM HEPES pH 7.3) and washed three times. The worm pellet was concentrated and resuspended in a solution containing 1 mg/ml Texas Red-dextran (40,000 MW, D1829, Molecular Probes). The samples were incubated for 90 min while shaking at 300 rpm in the dark. The dye in solution was removed by washing the samples with egg buffer until the solution was clear, and animals were either imaged directly or transferred to standard culture plates for 1 hour prior to imaging. Animals were paralyzed in 10 mM Tetramisole, transferred to a 5 % agarose pad on a glass slide, and imaged by spinning disk confocal microscopy.

### FRAP experiments and analysis

For FRAP assays, laser power was adjusted in each experiment to avoid complete photobleaching of the selected area. Photobleaching was performed on a circular region with a diameter of 30 or 40 px (respectively 3.33 or 4.44 µm) at the cortex, and recovery was followed at 5 second intervals for 15 minutes. Time-lapse movies were analyzed in ImageJ. The size of the area for FRAP analysis was defined by the full width at half-maximum of an intensity plot across the bleached region in the first post-bleach frame. For each time-lapse frame, the mean intensity value within the bleached region was determined, and the background, defined as the mean intensity of a non-bleached region outside the animal, was subtracted. The mean intensities within the bleached region were corrected for acquisition photobleaching per frame using the background-subtracted mean intensity of a similar non-bleached region at the cortex, which was normalized to the corresponding pre-bleach mean intensity. FRAP recovery was calculated as the change in corrected intensity values within the bleached region. The first frame after bleach was defined as 0, and the mean intensity of the 10 frames before bleach as 1. Curve fitting was done using GraphPad Prism 8, on averaged recovery data per sample using non-linear regression analysis (least squares regression). One and two-phase association were tested and, in all cases, data were best fitted with a two-phase curve.

### Feeding RNAi

RNAi clones for *bbln-1, ifb-2, ifc-2, ifd-2* and *let-413* were obtained from the genome wide Vidal full-length HT115 RNAi feeding library derived from the ORFeome 3.1 collection (Rual et al., 2004). The RNAi clone for *ifc-1* was obtained from the genome wide Ahringer fragment HT115 RNAi feeding library (Kamath et al., 2003), supplied through Source BioScience. All clones were verified using Sanger sequencing. RNAi clones for *ifo-1, ifd-1, ifp-1* and *sma-5* were generated by subcloning the corresponding cDNA into a modified L4440 RNAi feeding vector, containing a linker with AscI and NotI restriction sites. To specifically target *act-5*, and no other actin isoforms that share extensive sequence homology, we followed a previously described strategy and designed an RNAi clone against the unique 3’UTR (MacQueen et al., 2005). For all custom RNAi clones, fragments were amplified from a cDNA library, digested with AscI/NotI, ligated into AscI/NotI-digested L4440, and transformed into *E. coli* DH5α. Single colonies were isolated, plasmid DNA was purified, and presence of an insert was confirmed by Sanger sequencing. Correct clones were re-transformed into *E. coli* HT115, again confirmed by sequencing and stored at −80 °C in 50% glycerol (1:1). All primer pairs are listed in Table S2.

For feeding RNAi experiments, bacterial clones were pre-cultured in 2 ml Lysogeny Broth (LB) supplemented with 100 µg/ml ampicillin and 2.5 µg/ml tetracycline at 37 °C while rotating at 200 rpm for 6–8 hours, and then transferred to new tubes with a total volume of 10 mL and cultured overnight. An HT115 bacterial clone expressing the L4440 vector lacking an insert was used as a control in feeding experiments. To induce production of dsRNA, cultures were incubated for 90 min in the presence of 1 mM Isopropyl β-D-1-thiogalactopyranoside (IPTG). Bacterial cultures were pelleted by centrifugation at 3220 g for 15 min and concentrated 5x. NGM agar plates supplemented with 100 µg/ml ampicillin and 1 mM IPTG were seeded with 250 μl of bacterial suspension, and kept at room temperature for 48 hour in the dark. L1 or L4 hermaphrodites were placed on the seeded RNAi plates and incubated at 20 °C (Timmons and Fire, 1998).

### Electron microscopy

Young adult animals were transferred into a 100 µm deep membrane carrier containing 20% bovine serum albumin in M9 worm buffer (22 mM KH_2_PO_4_, 42 mM Na_2_HPO_4_, 86 mM NaCl, 1 mM MgSO_4_) and then high-pressure frozen in a Leica EM Pact high-pressure freezer. A minimum of five samples with 10-20 animals were frozen per experiment. Quick freeze substitution with agitation using 1% OsO_4_, 0.2% uranyl acetate in acetone followed by rapid epoxy resin embedding was performed as previously described (McDonald, 2014; Reipert et al., 2018). Subsequently, 50 nm thick sections of the embedded samples were prepared using a Leica UC7. These were contrasted for 10 min in 1% uranyl acetate in ethanol and Reynolds lead citrate and recorded at 100 kV on a Hitachi H-7600 transmission electron microscope (Tokyo, Japan).

### Auxin Inducible Degradation

Auxin treatment was performed by transferring worms to NGM plates seeded with *E. coli* OP50 and containing 1 mM auxin. To prepare plates, auxin (Alfa Aesar A10556) was diluted into the autoclaved NGM agar medium after cooling to 60 °C prior to plate pouring. Plates were kept for a maximum of 2 weeks in the dark at 4 °C. To obtain semi-synchronized worm populations, 40 adults were placed on NGM plates seeded with *E. coli* OP50 and allowed to lay eggs. After 1 hour of egg laying, plates were carefully washed with M9 (0.22 M KH2 PO4, 0.42 M Na2 HPO4, 0.85 M NaCl, 0.001 M MgSO4) buffer in order to remove larvae and adults but leave the eggs behind.

### Affinity-purification and mass spectrometry analysis

#### GFP pull-down from C. elegans

Animals endogenously expressing GFP-tagged BBLN-1 or control animals expressing an integrated GFP transgene (Waaijers et al., 2016) were grown on 6–8 9 cm NGM plates until starvation, to enrich for L1 animals. Animals were then transferred into 250 ml of S-Medium supplemented with 1% Penn/Strep (Life Technologies), 0.1% nystatin (Sigma) and OP50 bacteria obtained from the growth of a 400 ml culture. Animals were grown at 20 °C at low shaking for 96 hours and were harvested and cleaned using a sucrose gradient, as previously described (Waaijers et al., 2016) with one exception being the inclusion of MgSO_4_ in the M9 medium. Worms were distributed into 15 ml TPX tubes (Diagenode) to reach 200–400 µl worm pellet per tube, and were washed with lysis buffer (25mM Tris-HCl pH 7.5, 150mM NaCl, 1mM EDTA, 0.5% IGEPAL CA-630, 1X cOmplete Protease Inhibitor Cocktail (Roche)). The liquid was removed, and the sample was flash frozen in liquid nitrogen for storage at −80 °C.

To lyse the worms, tubes were thawed on ice and ice-cold lysis buffer was added to reach a total volume of 2 ml. Tubes were sonicated for 10 mins (sonication cycle: 30 sec ON, 30 sec OFF) at 4 °C in a Bioruptor ultrasonication bath (Diagenode) at high energy setting. After lysis, lysates were cleared by centrifugation and protein levels were measured using the Bradford BCA assay (Thermo Scientific).

Immunoprecipitation was performed using GFP-Trap Magnetic Agarose beads (Chromotek) according to manufacturer’s protocol, using 25 µl of beads per sample. To prep the beads, they were first equilibrated in wash buffer (10 mM Tris/Cl pH 7.5, 150 mM NaCl, 0.5 mM EDTA, 0.1% IGEPAL CA-630), blocked with 1% BSA for 1 hour, then washed 4 times with wash buffer. Next, lysate was added to the beads and they were incubated for 1 hour tumbling end-over-end. Lysate was then removed, and the beads were washed 4 times in wash buffer. After the final wash step, all liquid was removed, and the beads were flash frozen with liquid nitrogen. The experiment was performed in triplicate (biological replicates) and processed on independent days.

#### Biotin-streptavidin pull-down from cells

Confluent HEK293T cells were split in a 1:3 dilution 24 hours before transfection. Cells were transfected with overexpression constructs indicated in figure legend together with BirA. 1 μg/μL PEI (Polyethylenimine HCl MAX Linear MW 40000 (PolySciences, 24765–2)) and 1 μg/ μL DNA (3:1) were mixed in Ham’s F10 and incubated for 5 minutes at room temperature. The mixture was added to cells and incubated for 24 hours to allow expression. The cells were harvested in ice-cold PBS and lysed with lysis buffer (150 mM Tris-HCl pH 7.5, 150 mM NaCl, 1% Triton X-100 and cOmplete protease inhibitor cocktail (Roche)). 90% of each cell lysate was centrifuged at 13,000 rpm for 5 minutes and the supernatants were transferred and incubated with streptavidin beads (Dynabeads M-280, Invitrogen), which were already blocked by 0.2% Chicken Egg Albumine (Sigma). The remaining cell lysates were denatured with SDS/DTT sample buffer and used as input sample. Beads were incubated for 40 minutes at 4 °C, before washing 5 times with washing buffer (100mM Tris pH7.5, 150 mM NaCl, 0.5% Triton X-100 and 0.5x protease inhibitor cocktail).

#### Western blot analysis for bublin/keratin pull-down

For western blot assays, pull-down samples were eluted with SDS/DTT sample buffer and boiled for 5 min at 95 °C. Both pull-down and input were loaded into 10% SDS-PAGE gels and transferred to nitrocellulose membrane. Membranes were blocked with 2% BSA (bovine serum albumin) in PBS/0.05% Tween-20. Primary antibodies were diluted in blocking buffer and incubated with the membranes overnight at 4 °C, washed 3 times with PBS/0.05% Tween-20 and incubated with IRDye 680LT and IRDye 800CW antibodies (LI-COR Biosciences) for 45 min at RT. Membranes were washed 3 times with PBS/0.05% Tween-20 before imaging on an Odyssey Infrared Imaging system (LI-COR Biosciences).

#### Mass spectrometry analysis for BBLN-1/bublin

Streptavidin and anti-GFP beads after affinity purification were resuspended in 15 μl of 4× Laemmli sample buffer (Biorad), boiled at 99 °C for 10 min and supernatants were loaded on 4–12% Criterion XT Bis–Tris precast gel (Biorad). The gel was fixed with 40% Methanol and 10% acetic acid and then stained for 1 hour using colloidal coomassie dye G-250 (Gel Code Blue Stain, Thermo Scientific). Each lane from the gel was cut and placed in 1.5 ml tubes. Samples were then washed with 250 μl of water, followed by 15 min dehydration in acetonitrile. Proteins were reduced (10mM DTT, 1 hour at 56 °C), dehydrated and alkylated (55mM iodoacetamide, 1 hour in the dark). After two rounds of dehydration, trypsin was added to the samples and incubated overnight at 37 °C. Peptides were extracted with acetonitrile, dried down and reconstituted in 10% formic acid prior to MS analysis.

Samples were analyzed on an Orbitrap Q-Exactive mass spectrometer (Thermo Fisher Scientific) coupled to an Agilent 1290 Infinity LC (Agilent Technologies). Peptides were loaded onto a trap column (Reprosil pur C18, Dr. Maisch, 100 μm × 2 cm, 3 μm; constructed in-house) with solvent A (0.1% formic acid in water) at a maximum pressure of 800 bar and chromatographically separated over the analytical column (Poroshell 120 EC C18, Agilent Technologies, 100 μm x 50 cm, 2.7 μm) using 90 min linear gradient from 7% to 30% solvent B (0.1% formic acid in acetonitrile) at a flow rate of 150 nl min^−1^. The mass spectrometers were used in a data-dependent mode, which automatically switched between MS and MS/MS. After a survey scan from 375 to 1600m/z the 10 most abundant peptides were subjected to HCD fragmentation. MS spectra were acquired with a resolution > 30,000, whereas MS2 with a resolution > 17,500.

Raw data files were converted to mgf files using Proteome Discoverer 1.4 software (Thermo Fisher Scientific). Database search was performed using the *C. elegans* or the human database and Mascot (version 2.5.1, Matrix Science, UK) as the search engine. Carbamidomethylation of cysteines was set as a fixed modification and oxidation of methionine was set as a variable modification. Trypsin was set as cleavage specificity, allowing a maximum of two missed cleavages. Data filtering was performed using a percolator, resulting in 1% false discovery rate (FDR). Additional filters were search engine rank 1 and mascot ion score > 20.

Crapome (Mellacheruvu et al., 2013) was used to analyze BBLN-1 interacting proteins in three biological replicas and bublin binding proteins in a single experiment, using proteins identified in the GFP pull downs as control. Significance analysis of interactome (SAINT) score (Choi et al., 2011) and simpler fold-change (FC) calculations FC-A and FC-B were derived from the Crapome analysis by averaging the spectral counts across the controls. FC-A averages the counts across all controls while the more stringent FC-B takes the average of the top 3 highest spectral counts for the abundance estimate.

### Antibodies and Immunofluorescence Cell Staining

We used rabbit polyclonal antibodies against bublin (Sigma-Aldrich Cat# HPA020725, RRID:AB_1845816), mouse monoclonal antibodies against PCNT (Abcam Cat# ab28144, RRID:AB_2160664), and rat monoclonal antibodies against tyrosinated α-tubulin (YL1/2; Thermo Fisher Scientific Cat# MA1-80017, RRID:AB_2210201) and KRT8 (DSHB Cat# TROMA-I, RRID:AB_531826). We used Alexa Fluor 568-conjugated Phalloidin to stain actin (Life Technologies). We used Alexa Fluor 488-, 594- and 647-conjugated goat antibodies against respectively mouse, rabbit and rat (Life Technologies) as secondary antibodies.

HeLa cells were fixed with −20 °C methanol or 4% PFA for 10 min and permeabilized with 0.1% Triton X-100 in phosphate-buffered saline (PBS) for 10 min. Subsequent washing and labeling steps were carried out in PBS supplemented with 2% bovine serum albumin and 0.05% Tween-20. Slides were rinsed in 70 and 100% ethanol and mounted in Vectashield mounting medium supplemented with DAPI (Vector Laboratories).

Mouse small intestinal organoids were fixed in suspension using ice-cold 4% PFA (Aurion) and immunolabeled as described previously (van Ineveld et al., 2020).

### Statistical analysis

All statistical analyses were performed using GraphPad Prism 8. For population comparisons, a D’Agostino & Pearson test of normality was first performed to determine if the data was sampled from a Gaussian distribution. For data drawn from a Gaussian distribution, comparisons between two populations were done using an unpaired t-test, with Welch’s correction if the SDs of the populations differ significantly, and comparisons between >2 populations were done using a one-way ANOVA if the SDs of the populations differ significantly. For data not drawn from a Gaussian distribution, a non-parametric test was used (Mann-Whitney for 2 populations and Kruskal-Wallis for >2 populations). ANOVA and non-parametric tests were followed up with multiple comparison tests of significance (Dunnett’s, Tukey’s, Dunnett’s T3 or Dunn’s). Tests of significance used and sample sizes are indicated in the figure legends. No statistical method was used to pre-determine sample sizes. No samples or animals were excluded from analysis. The experiments were not randomized, and the investigators were not blinded to allocation during experiments and outcome assessment.

## Notes

### Competing Interest Statement

The authors have declared no competing interest.

### Summary of Updates

This version changes the names of the proteins BUBL-1 and C9orf16 to the newly approved names BBLN-1 and bublin (BBLN), respectively.

