## Supplemental information for "BBLN-1 is essential for intermediate filament organization and apical membrane morphology"

### Figure S1

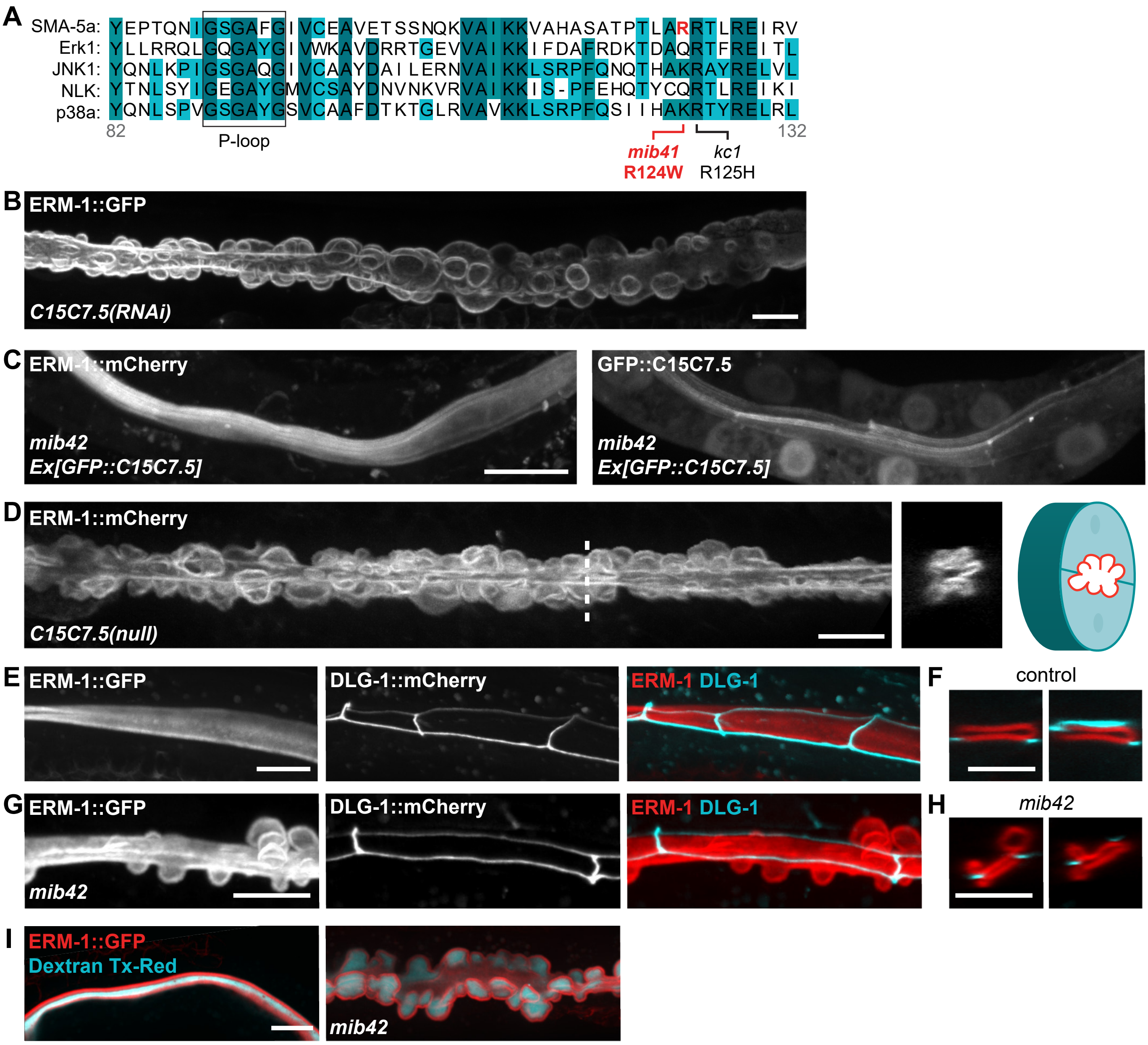


**Figure S1 –** (**A**) Partial sequence alignment of the kinase domain of SMA-5 and human MAPK family kinases. The boxed region corresponds to the conserved phosphate-binding loop (P-loop). The positions of *mib41* (red) and *kc1* (black) mutations are indicated below. (**B**) L3 *C15C7.5(RNAi)* larva expressing an erm-1::GFP knock-in. (**C**) mib42 L3 larva expressing an erm-1::mCherry knock-in and a GFP::C15C7.5 transgene driven by the intestine-specific *vha-6* promoter. (**D**) Intestinal apical membrane morphology in a *C15C7.5(null)* L4 animal visualized with an endogenous ERM-1::mCherry reporter. Dotted line in left panel indicates position of cross-section view. Schematic depicts cross-section with apical membrane in red. (**E–H**) Organization of the C. elegans apical junctions (CeAJs) in intestinal cells of control (E, F) and *mib42* mutant animals (G, H), visualized by an endogenous DLG-1::mCherry reporter in larvae in which the apical membrane is labelled by ERM-1::GFP. (**I**) Intestinal epithelium integrity assessed by feeding membrane-impermeable Dextran conjugated to a Texas-Red fluorescent dye in animals expressing ERM-1::GFP.

#### Figure S2

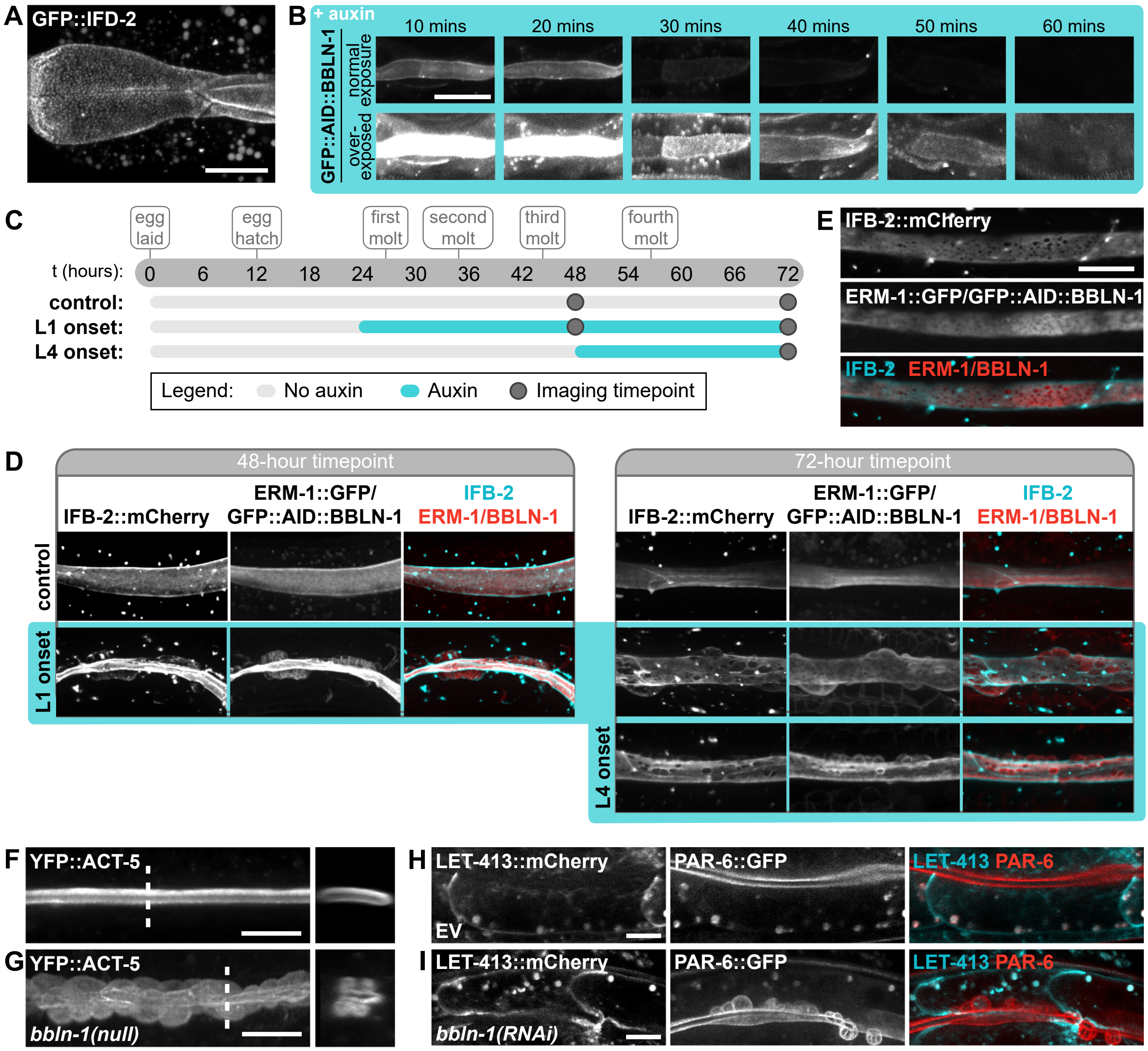


**Figure S2 –** (**A**) Localization of GFP::IFD-2 in the anterior most section of the intestine, highlighting the punctate localization pattern of IFD-2. (**B**) Distribution of GFP::AID::BBLN-1 in L1 larvae in the presence of 1 mM auxin for 10 to 60 mins. Degradation was induced in the intestine. Bottom row depicts the same images as top row, but computationally overexposed in ImageJ (Fiji) to visualize low levels of fluorescence. (**C**) Schematic overview of experiment design for the data in D. Eggs were laid on t = 0 and allowed to develop on NGM plates for 24 (L1 onset) or 48 (L4 onset) hours before being transferred to plates containing 1 mM auxin. Larvae were imaged after 24 (L1 and L4 onset, 48-hour timepoint) and 48 (L1 onset, 72-hour timepoint) hours of degradation. (**D**) Distribution of IFB-2::mCherry, ERM-1::GFP and GFP::AID::BBLN-1 upon intestine specific degradation of GFP::AID::BBLN-1 in L4 larvae (left, 48-hour timepoint) and young adults (right, 72-hour timepoint). (**E**) L4 larva displaying 'holes' in IFB-2::mCherry expression pattern upon 24 hours of degradation from the L1 stage. (**F, G**) Organization of the YFP::ACT-5 transgene in intestinal cells as seen in lateral and cross-section views as indicated by the dashed lines. (**H, I**) Apicobasal polarity of the intestine visualized by an LET-413::mCherry endogenous reporter for the basolateral domain and an endogenous PAR-6::GFP marker for the apical domain.

#### Figure S3

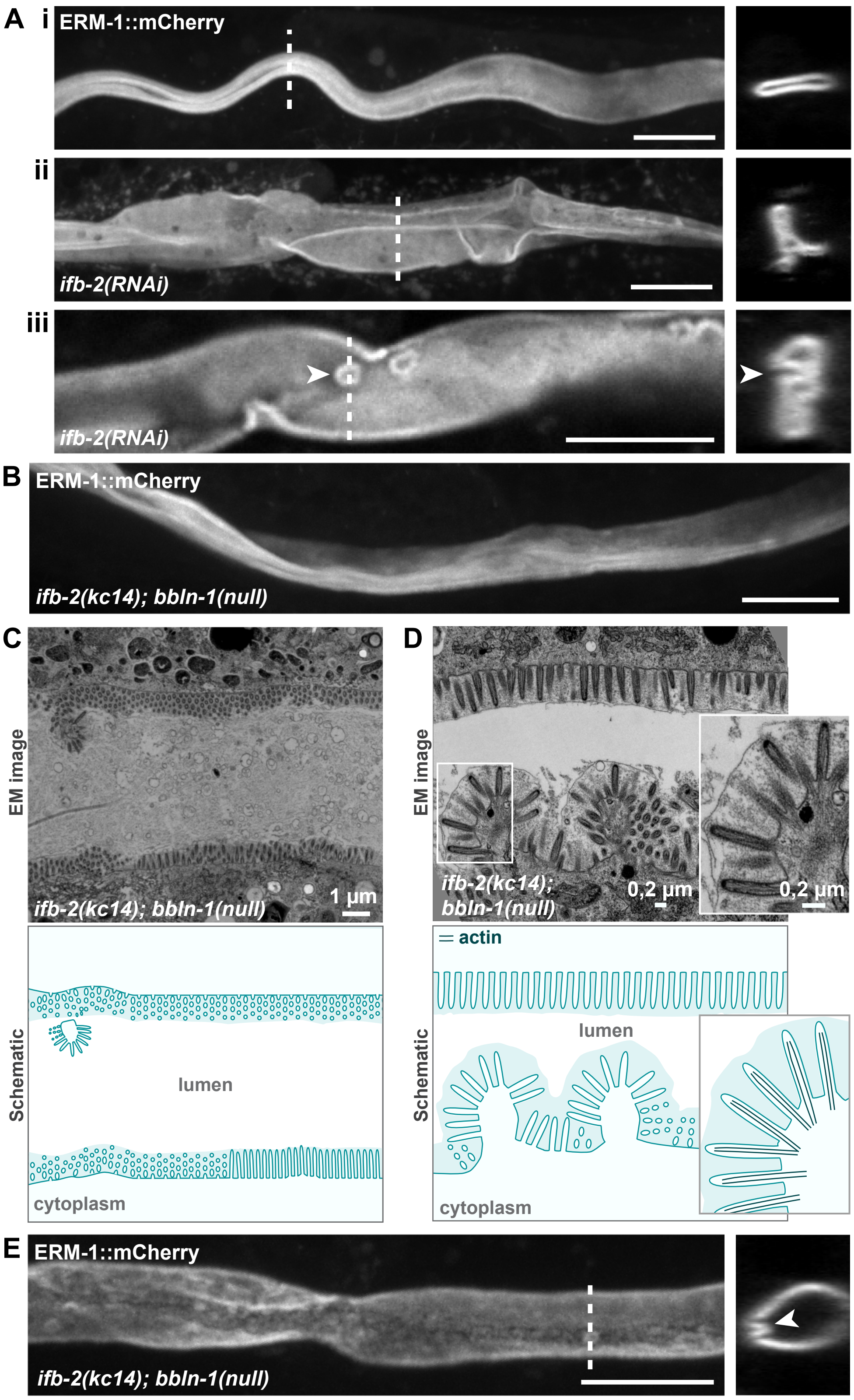


**Figure S3 –** (**A**) Apical membrane morphology in intestinal cells visualized by ERM-1::mCherry. Dashed lines indicate the position of the cross-sections. Arrowheads in (iii) point to apical membrane protrusion towards the lumen (same protrusion is indicated in lateral and cross-section views). (**B**) Apical membrane morphology visualized by ERM-1::mCherry in *ifb-2; bbln-1* double knock-out. (**C, D**) Ultrastructure of the apical domain in intestinal cells visualized by transmission electron microscopy. Boxed region in (D) is shown in zoom-in. Schematics indicate actin bundles in dark blue in the zoom inset. (**E**) Apical membrane morphology visualized by ERM-1::mCherry in *ifb-2; bbln-1* double knockout. Arrowhead indicates apical membrane protrusion towards the lumen.

#### Figure S4

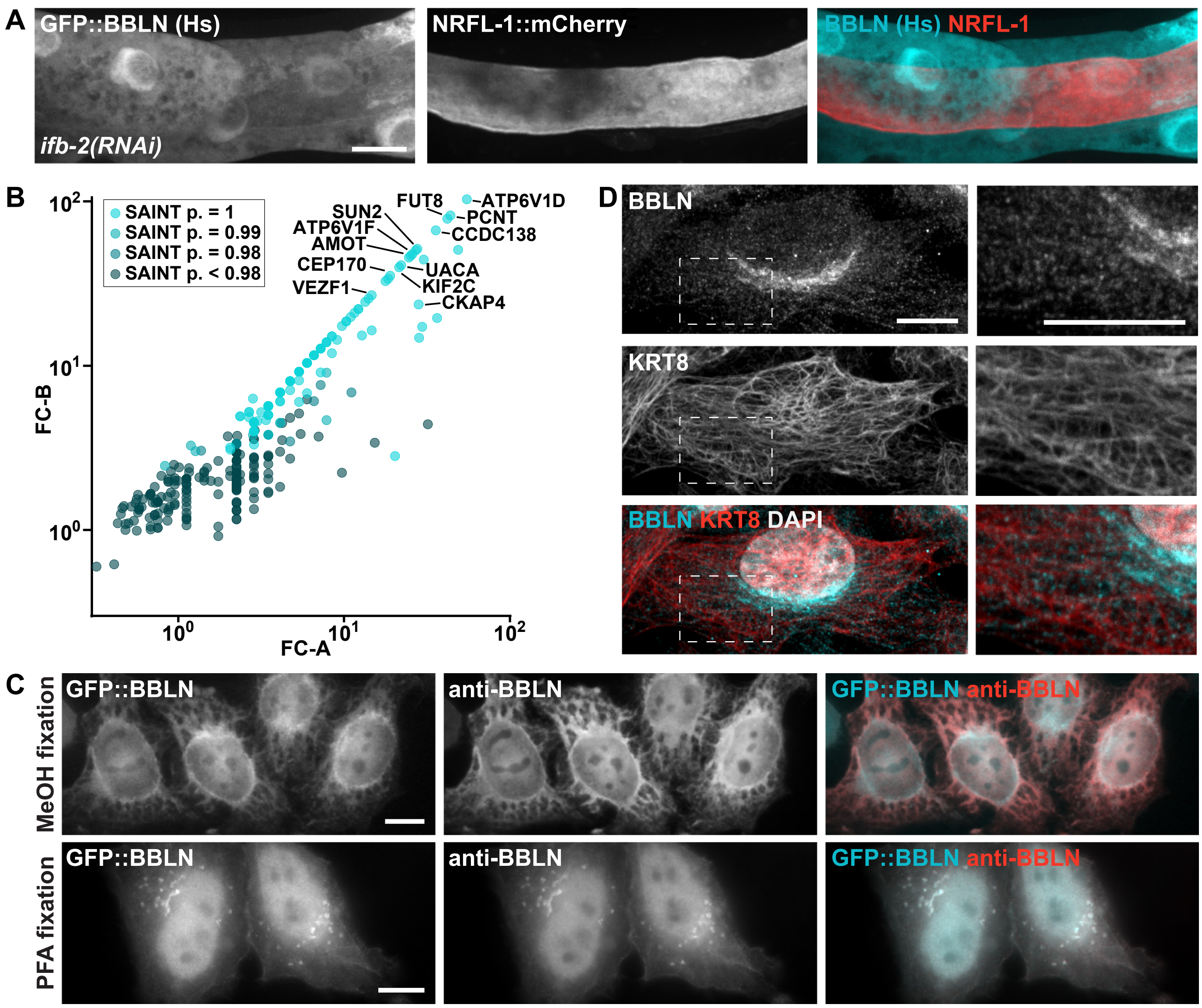


**Figure S4 –** (**A**) Larva expressing intestinal GFP-tagged bublin (BBLN (Hs)) fed with a bacterial RNAi clone against *ifb-2*. An endogenous NRFL-1::mCherry fusion was used as an apical membrane marker. (**B**) Mass spectrometry hits for bublin plotted as correlation between fold-change (FC) score A and more stringent FC score B. Data points are color coded for different SAINT probability scores. (**C**) Upright fluorescence microscopy images of HeLa cells transfected with GFP-tagged bublin (BBLN (Hs)) and stained with bublin (BBLN) antibody fixed with either -20 °C methanol (top) or 4% PFA (bottom). (**D**) Confocal images of HeLa cells stained with bublin (BBLN) antibody, keratin 8 (KRT8) antibody and DAPI and fixed by 4% PFA. Boxed region indicates location of zoom-in on the right.

**Table S1 – List of strains used**

| Strain | Genotype |
|---|---|
| N2 | wild type |
| BOX64 | *mibIs39[Prps-27::GFP-2xTEV-Avi 10 ng/ul + Prab-3::mCherry 5 ng/ul + lambda DNA 65 ng/ul] I* |
| BOX213 | *erm-1(mib15[erm-1::eGFP]) I* |
| BOX320 | *erm-1(mib15[erm-1::eGFP]) I; sma-5a(mib41[C370T]) X* |
| BOX321 | *erm-1(mib15[erm-1::eGFP]) I; bbln-1(mib42[C13T]) X* |
| BOX414 | *bbln-1(mib71[eGFP::bbln-1]) X* |
| BOX459 | *bbln-1(mib79[bbln-1::mkate2(co)]) X* |
| BOX415 | *erm-1(mib40[erm-1::AID::mCherry]) I; bbln-1(mib70[Pbbln-1::eGFP1-3, X:3151104..3153328]) X* |
| BOX427 | *erm-1(mib40[erm-1::AID::mCherry]) I; bbln-1(mib71[eGFP::bbln-1]) X* |
| BOX435 | *ifb-2(mib74[ifb-2::mCherry]) II; bbln-1(mib70[Pbbln-1::eGFP1-3, X:3151104..3153328]) X* |
| BOX436 | *ifb-2(mib74[ifb-2::mCherry]) II; bbln-1(mib71[eGFP::bbln-1]) X* |
| BOX514 | *erm-1(mib15[erm-1::eGFP]) I; ifb-2(mib74[ifb-2::mCherry]) II* |
| BOX515 | *erm-1(mib15[erm-1::eGFP]) I; ifb-2(mib74[ifb-2::mCherry]) II; bbln-1(mib70[Pbbln-1::eGFP1-3, X:3151104..3153328]) X* |
| BOX303 | *erm-1(mib40[erm-1::AID::mCherry]) I* |
| BOX330 | *erm-1(mib40[erm-1::AID::mCherry]) I; bbln-1(mib42[C13T]) X* |
| BOX307 | *erm-1(mib15[erm-1::eGFP]) I; bbln-1(mib42[Q5STOP]) X; dlg-1(mib23[dlg-1::mCherry-LoxP]) X* |
| BOX368 | *erm-1(mib15[erm-1::eGFP]) I; dlg-1(mib23[dlg-1::mCherry-LoxP]) X* |
| BOX454 | *ifb-2(mib74[ifb-2::mCherry]) II; bbln-1(mib70[Pbbln-1::eGFP1-3, X:3151104..3153328]) X; dlg-1(mi-b35[dlg-1::AID::eGFP-LoxP]) X* |
| BOX455 | *ifb-2(mib74[ifb-2::mCherry]) II; dlg-1(mib35[dlg-1::AID::eGFP-LoxP]) X* |
| BOX456 | *erm-1(mib40[erm-1::AID::mCherry])I; ifc-2a::yfp(kc16)X* |
| BOX457 | *erm-1(mib40[erm-1::AID::mCherry]) I; ifc-2a::yfp(kc16) X; bbln-1(mib70[Pbbln-1::eGFP1-3, X:3151104..3153328]) X* |
| BOX614 | *erm-1(mib40[erm-1::AID::mCherry]) I; ifd-2(mib94[eGFP::ifd-2]) X* |
| BOX615 | *erm-1(mib40[erm-1::AID::mCherry]) I; ifd-2(mib94[eGFP::ifd-2]) X; bbln-1(mib70[Pbbln-1::eGFP1-3, X:3151104..3153328]) X* |
| JM125 | *Is[Pges-1::YFP::ACT-5]* |
| BOX438 | *bbln-1(mib70[Pbbln-1::eGFP1-3, X:3151104..3153328]) X; Is[Pges-1::YFP::ACT-5]* |
| BOX251 | *par-6(mib24[par-6::eGFP-LoxP]) I; let-413(mib29[let-413::mCherry-LoxP]) V* |
| BOX544 | *gip-2(lt19[gip-2::GFP]::loxP::cb-unc-119(+)::loxP) I; bbln-1(mib93[mCherry::bbln-1]) X* |
| BOX632 | *mibIs48[Pelt-2::TIR-1::tagBFP2-Lox511::tbb-2-3'UTR, IV:5014740-5014802 (cxTi10882 site)]) IV; bbln--1(mib111[eGFP::AID::bbln-1]) X* |
| BOX637 | *erm-1(mib15)I; IFB-2(mib74[IFB-2::mCherry]) II; mibIs48[Pelt-2::TIR-1::tagBFP2-Lox511::tbb-2-3'UTR, IV:5014740-5014802 (cxTi10882 site)]) IV; bbln-1(mib111[GFP::AID::BBLN-1]) X* |
| BJ364 | *erm-1(mib40[erm-1::AID::mCherry]) I; ifb-2(kc14) II; bbln-1(mib70[Pbbln-1::eGFP1-3, X:3151104..3153328]) X* |
| OD2509 | *gip-2(lt19[gip-2::GFP]::loxP::cb-unc-119(+)::loxP)I; unc-119(ed3)III* |

**Table S2 – List of DNA reagents used**

| | | |
|---|---|---|
| **Reagents to generate RNAi clones** | | |
| *sma-5* | | |
| | Forward | aggcgcgccACATTGTCCCTCTCCGTGAC |
| | Reverse | agcggccgcTTCGTCGTCATGCTTCTTTG |
| *ifp-1* | | |
| | Forward | aggcgcgccTGACCACCATAGCCGAACTT |
| | Reverse | agcggccgcTTTGAAGCCACCAACGTCTG |
| *ifd-1* | | |
| | Forward | aggcgcgccTCAAAACCGGGTTCTCGAGA |
| | Reverse | agcggccgcTTCACTGCGGAGGTTGATCT |
| *ifo-1* | | |
| | Forward | aggcgcgccCCTACAAGTCGACTTGAATGCAGC |
| | Reverse | agcggccgcAGTGAAGTGGGCGAGTGATG |
| *act-5* | | |
| | Forward | aggcgcgccaacatgtgccttccatttctaggcg |
| | Reverse | agcggccgcAGAAAATGAAGTATCTCATGGAATTTG |

| | | |
|---|---|---|
| **Reagents to generate *erm-1::mCherry*** | | |
| sgRNA 1 | | |
| | Forward | tcttAAGACTCTCCGTCAAATCCG |
| | Reverse | aaacCGGATTTGACGGAGAGTCTT |
| sgRNA 2 | | |
| | Forward | tcttACTCTCCGTCAAATCCGTGG |
| | Reverse | aaacCCACGGATTTGACGGAGAGT |
| Repair template primers | | |
| | LH arm forward | GGCTGCTCTTCgTGGGGAGgttcgtattttaaaaaaactcg |
| | LH arm reverse 1 | TGATCGATTCTTCGTTTTGTGTTTCCTCCcCtaATcTGcCGGAGAGTCTTGTACTTGTCG |
| | LH arm reverse 2 | GGGTGCTCTTCgCGCCATATTTTCGTATTGATCGATTCTTCGTTTTGTGTTTCC |
| | RH arm forward | GGCTGCTCTTCgACGTAAttatttgttctatcgtatttccttt |
| | RH arm reverse | GGGTGCTCTTCgTACgctccatcgaaacccttgga |
| Genotyping primers | | |
| | Forward | CTGTCACTGACTACGACGTTCTG |
| | Reverse | CCCGAGGAGAAGCACACATG |

| | | |
|---|---|---|
| **Reagents to generate *bbln-1(mib70)*** | | |
| crRNA | | |
| | N-terminal | ctcatttcagttgaacacaa |
| | C-terminal | gaagagagttcaatctgtca |
| Repair template ssODN | | cgtctttttctccatttcctcatttcagttgaacacaatgTCCAAGGGAGAGGAGCTCTTCACCGGAGTCGTC-CCAATCCTCGTCGAGCTCGACGGAGTCAAGGAGTTCGTCACCGCTGCCGGAATCACCCACG-GAATGGACGAGCTCTACAAGtaagagagttcaatctgtcaaacgcaccgaaaaataaa |
| Genotyping primers | | |
| | Forward | atcatcacccatctccaacc |

| | |
|---|---|
| Reverse | CGCGCATCTTGACAATAGGC |

| **Reagents to generate *gfp::bbln-1*** | |
|---|---|
| crRNA | |
| Step 1 | ctcatttcagttgaacacaa |
| Step 2 | cgtcgagctcgacggagtca |
| Repair template ssODN | |
| Step 1 | cgtctttttctccatttcctcatttcagttgaacacaatgTCCAAGGGAGAGGAGCTCTTCACCGGAGTCGTC-<br>CCAATCCTCGTCGAGCTCGACGGAGTCAAGGAGTTCGTCACCGCTGCCGGAATCACCCACG-<br>GAATGGACGAGCTCTACAAGgtcgttgagcagaaagagcaagagcctattgtcaa |
| Repair template primers | |
| Step 2 forward | CCAAGGGAGAGGAGCTCTTCA |
| Step 2 reverse | CTTGTAGAGCTCGTCCATTC |
| Repair template PCR | CCAAGGGAGAGGAGCTCTTCACCGGAGTCGTCCCAATCCTCGTCGAGCTCGACGGAGACGTCA-<br>ACGGACACAAGTTCTCCGTCTCAGGAGAGGGAGAGGGAGACGCCACCTACGGAAAGCTCACCC-<br>TCAAGTTCATCTGCACCACCGGAAAGCTCCCAGTCCCATGGCCAACCCTCGTCACCACCTTCACT-<br>TACGGAGTCCAATGCTTCTCCCGTTACCCAGACCACATGAAGCGTCACGACTTCTTCAAGTCCGC-<br>CATGCCAGAGGGATACGTCCAAGAGCGTACCATCTTCTTCAAGgtaagtttaaacattaattaatactaac-<br>taaccctgattatttaaattttcagGACGACGGAAACTACAAGACCCGTGCCGAGGTCAAGTTCGAGGGA-<br>GACACCCTCGTCAACCGTATCGAGCTCAAGgtaagtttaaacagttcggtactaactaaccatacatatttaaattt-<br>tcagGGAATCGACTTCAAGGAGGACGGAAACATCCTCGGACACAAGCTCGAATACAACTACA-<br>ACTCCCACAACGTCTACATCATGGCCGACAAGCAAAAGAACGGAATCAAGGTCAACTTCAAGgt-<br>aagtttaaacatgattttactaactaactaatctgatttaaattttcagATCCGTCACAACATCGAGGACGGATCTG-<br>TCCAACTCGCCGACCACTACCAACAAAACACCCCAATCGGAGACGGACCAGTCCTCCTCCCA-<br>GACAACCACTACCTCTCCACCCAATCCGCCCTCTCCAAGGACCCAAACGAGAAGCGTGACCA-<br>CATGGTCCTCAAGGAGTTCGTCACCGCTGCCGGAATCACCCACGGAATGGACGAGCTCTACAAG |
| Genotyping primers | |
| Forward | ATCATCACCCATCTCCAACC |
| Reverse | CGCGCATCTTGACAATAGGC |

| **Reagents to generate *bbln-1*::mKate2** | |
|---|---|
| crRNA | gtaccaattgaaaagcattc |
| Repair template 5'SP9 modified primers | |
| Forward | TGAAGAGAGTTCAATCTGTCAAACGCACCGAAAAAATGTCCGAGCTCATCAAGGAG |
| Reverse | gtgtacatgtaccaattgaaaagcattctggtTTAACGGTGTCCGAGCTTGGAT |
| Repair template PCR | ATGTCCGAGCTCATCAAGGAGAACATGCACATGAAGCTCTACATGGAGGGAACCGTCAACAAC-<br>CACCACTTCAAGTGCACCTCCGAGGGAGAGGGAAAGCCATACGAGGGAACCCAAACCATGCG-<br>TATCAAGgtaagtttaaacatatatatactaactaaccctgattatttaaattttcagGCCGTCGAGGGAGGACCAC-<br>TCCCATTCGCCTTCGACATCCTCGCCACCTCCTTCATGTACGGATCCAAGACCTTCATCAACCA-<br>CACCCAAGGAATCCCAGACTTCTTCAAGCAATCCTTCCCAGAGGGATTCACCTGGGAGCGTGT-<br>CACCACCTACGAGGACGGAGGAGTCCTCACCGCCACCCAAGACACCTCCCTCCAAGACGGAT-<br>GCCTCATCTACAACGTCAAGgtaagtttaaacagttcggtactaactaaccatacatatttaaattttcagATCCGTG-<br>GAGTCAACTTCCCATCCAACGGACCAGTCATGCAAAAGAAGACCCTCGGATGGGAGGCCTCCAC-<br>CGAGACCCTCTACCCAGCCGACGGAGGACTCGAGGGACGTGCCGACATGGCCCTCAAGCTCG-<br>TCGGAGGAGGACACCTCATCTGCAACCTCAAGgtaagtttaaacatgattttactaactaactaatctgattta-<br>aattttcagACCACCTACCGTTCCAAGAAGCCAGCCAAGAACCTCAAGATGCCAGGAGTCTACTACG-<br>TCGACCGTCGTCTCGAGCGTATCAAGGAGGCCGACAAGGAGACCTACGTCGAGCAACACGAGG-<br>TCGCCGTCGCCCGTTACTGCGACCTCCCATCCAAGCTCGGACACCGT |
| Genotyping primers | |
| Forward | GCAAATGAGCAGCATCCTG |

| | |
|---|---|
| Reverse | cgaggaaccaaatcattttcc |

| **Reagents to generate *ifb-2::mCherry*** | |
|---|---|
| crRNA | |
| Step 1 | gatgatggagatttcTTAAC |
| Step 2 | GTTCATGCGTTTCAAGGCCG |
| Repair template ssODN | |
| Step 1 | tcatcgaaaaagaaatcgattagatgatggagatttcTTACTTGTAGAGCTCGTCCATTCCTCCGGTGGAGT-<br>GACGTCCCTCGGCCTTGAAACGCATGAACTCCTTGATGATGGCCATGTTGTCCTCCTCTCCCTTG-<br>GAACGGGAAGAAGCGACCGTCGTCTGGATGTGCGAAGCCTTC |
| Repair template primers | |
| Step 2 forward | TCCAAGGGAGAGGAGGACAA |
| Step 2 reverse | CTTGTAGAGCTCGTCCATTC |
| Repair template PCR | TCCAAGGGAGAGGAGGACAACATGGCCATCATCAAGGAGTTCATGCGTTTCAAGGTCCACATG-<br>GAGGGATCAGTCAACGGACACGAGTTCGAGATCGAGGGAGAGGGAGAGGGACGTCCATAC-<br>GAGGGAACCCAAACCGCCAAGCTCAAGgtaagtttaaacatatatatactaactaaccctgattatttaaatttt-<br>cagGTCACCAAGGGAGGACCACTCCCATTCGCCTGGGACATCCTCTCCCCACAATTCATGTACG-<br>GATCAAAGGCCTACGTCAAGCACCCAGCCGACATCCCAGACTACCTCAAGCTCTCCTTCCCA-<br>GAGGGATTCAAGTGGGAGCGTGTCATGAACTTCGAGGACGGAGGAGTCGTCACCGTCACCCA-<br>AGACTCCTCCCTCCAAGACGGAGAGTTCATCTACAAGgtaagtttaaacagttcggtactaactaaccataca-<br>tatttaaattttcagGTCAAGCTCCGTGGAACCAACTTCCCATCCGACGGACCAGTCATGCAAAAGA-<br>AGACCATGGGATGGGAGGCCTCCTCCGAGCGTATGTACCCAGAGGACGGAGCCCTCAAGGGA-<br>GAGATCAAGCAACGTCTCAAGCTCAAGGACGGAGGACACTACGACGCCGAGGTCAAGACCACC-<br>TACAAGGCCAAGAAGCCAGTCCAACTCCCAGgtaagtttaaacatgattttactaactaactaatctgattta-<br>aattttcagGAGCCTACAACGTCAACATCAAGCTCGACATCACCTCCCACAACGAGGACTACAC-<br>CATCGTCGAGCAATACGAGCGTGCCGAGGGACGTCACTCCACCGGAGGAATGGACGAGCTCTA-<br>CAAG |
| Genotyping primers | |
| Forward | tcggtagctataaccgcttca |
| Reverse | caaggaaaggattcaatgggc |

| **Reagents to generate *Pvha-6::gfp::bbln-1::tbb-2 3'UTR* construct** | |
|---|---|
| *vha-6* promoter | |
| Forward | ctGCTCTTCgTGGTTGCCAGTGATGAATCCAAGCAC |
| Reverse | ctGCTCTTCgCATtttttatgggttttggtaggttttagtcg |
| *tbb-2* 3'UTR | |
| Forward | ctGCTCTTCGACGTAAgataaatgcaaaatcctttcaag |
| Reverse | ctGCTCTTCgTACtgagactttttcttggcggc |
| *bbln-1* gBlock | gcatggctGCTCTTCgAAGATGGTCGTTGAGCAGAAAGAGCAAGAGCCTATTGTCAAGATG-<br>CGCGACCGCAATGTCAACGCTGCTGCACATTCTGCGTTGGCTCGTGGAATTGAGGCACT-<br>CAACGAAGGAGAAGTGACCGAGGAGACGGAAGgtgaaaactcttctcagattcagattacttatagcattggt-<br>tttttcagAAATTCGCAAGCTGGACACCCAGCTTGATCATCTTAATGACTACATGTCTAAGATGGAT-<br>GAGCGTCTGAAGGCACACAACGACAGAATGATGGAGACGTTGAAGCAGCAGAAGGATGAGCG-<br>CGAAAAGAGACGTCGCAGCTTCCACGAGCGTATGTCCCAAAATCAATCTGAAGATGAGGAGT-<br>TCAAAAAGCAAATGAGCAGCATCCTGAAGAGAGTTCAATCTGTCAAACGCACCGAAAAAGGTc-<br>GAAGAGCagccgga |

| | |
|---|---|
| **Reagents to generate *Pvha-6::gfp::BBLN (Hs)::tbb-2 3’UTR* construct** | |
| *vha-6* promoter | |
| Forward | ctGCTCTTCgTGGTTGCCAGTGATGAATCCAAGCAC |
| Reverse | ctGCTCTTCgCATtttttatgggttttggtaggttttagtcg |
| *tbb-2* 3’UTR | |
| Forward | ctGCTCTTCGACGTAAgataaatgcaaaatcctttcaag |
| Reverse | ctGCTCTTCgTACtgagactttttcttggcggc |
| *BBLN (Hs)* gBlock | gcatggctGCTCTTCgAAGATGTCCGGACCAAACGGAGACCTCGGAATGCCAGTCGAGGCCGGA-GCCGAGGGAGAGGAGGACGGATTCGGAGAGGCCGAGTACGCCGCCATCAACTCCATGCTC-GACCAAATCAACTCCTGCCTCGACCACCTCGAGGAGAAGgtaagtttaaacatatatatactaactaaccc-tgattatttaaattttcagAACGACCACCTCCACGCCCGTCTCCAAGAGCTCCTCGAGTCCAACCGTCA-AACCCGTCTCGAGTTCCAACAACAACTCGGAGAGGCCCCATCCGACGCCTCCCCAGGTcGAAGA-GCagccggat |

| | |
|---|---|
| **Reagents to generate *gfp::ifd-2*** | |
| crRNA (5’) | TGGGTTGAGAGGGTCAGTCA |
| Repair template primers | |
| Forward | TATTCAAAACTAATTTCTAGAATAAAAACGCCATGTCCAAGGGAGAGGAGCTCTT |
| Reverse | ATGATTTTGCAGACGCGTTGGGTTGAGAGGGTCAGTCTTGTAGAGCTCGTCCATTC |
| Genotyping primers | |
| Forward | GGAACGGCTCAGTTTTTCTC |
| Reverse | CTACATATCGTGCCAATCGG |

| | |
|---|---|
| **Reagents to generate *mCherry::bbln-1*** | |
| crRNA | CTCATTTCAGTTGAACACAA |
| Repair template primers | |
| Forward | CGTCTTTTTCTCCATTTCCTCATTTCAGTTGAACACAATGTCCAAGGGAGAGGAGGACAA |
| Reverse | TTGACAATAGGCTCTTGCTCTTTCTGCTCAACGACCTTGTAGAGCTCGTCCATTC |
| Repair template PCR | TCCAAGGGAGAGGAGGACAACATGGCCATCATCAAGGAGTTCATGCGTTTCAAGGTCCACATG-GAGGGATCAGTCAACGGACACGAGTTCGAGATCGAGGGAGAGGGAGAGGGACGTCCATAC-GAGGGAACCCAAACCGCCAAGCTCAAGgtaagtttaaacatatatatactaactaaccctgattatttaaatttt-cagGTCACCAAGGGAGGACCACTCCCATTCGCCTGGGACATCCTCTCCCCACAATTCATGTACG-GATCAAAGGCCTACGTCAAGCACCCAGCCGACATCCCAGACTACCTCAAGCTCTCCTTCCCA-GAGGGATTCAAGTGGGAGCGTGTCATGAACTTCGAGGACGGAGGAGTCGTCACCGTCACCCA-AGACTCCTCCCTCCAAGACGGAGAGTTCATCTACAAGgtaagtttaaacagttcggtactaactaaccataca-tatttaaattttcagGTCAAGCTCCGTGGAACCAACTTCCCATCCGACGGACCAGTCATGCAAAAGA-AGACCATGGGATGGGAGGCCTCCTCCGAGCGTATGTACCCAGAGGACGGAGCCCTCAAGGGA-GAGATCAAGCAACGTCTCAAGCTCAAGGACGGAGGACACTACGACGCCGAGGTCAAGACCACC-TACAAGGCCAAGAAGCCAGTCCAACTCCCAGgtaagtttaaacatgattttactaactaactaatctgattta-aattttcagGAGCCTACAACGTCAACATCAAGCTCGACATCACCTCCCACAACGAGGACTACAC-CATCGTCGAGCAATACGAGCGTGCCGAGGGACGTCACTCCACCGGAGGAATGGACGAGCTCTA-CAAG |
| Genotyping primers | |
| Forward | ATCATCACCCATCTCCAACC |
| Reverse | CGCGCATCTTGACAATAGGC |

| **Reagents to generate *gfp::aid::bbln-1*** | |
|---|---|
| crRNA | CTCATTTCAGTTGAACACAA |
| Repair template 5’SP9 modified primers | |
| Forward | CGTCTTTTTCTCCATTTCCTCATTTCAGTTGAACACAATGTCCAAGGGAGAGGAGCTCTT |
| Reverse | TTGACAATAGGCTCTTGCTCTTTCTGCTCAACGACCTTCACGAACGCCGCCGCCT |
| Repair template PCR | cgtctttttctccatttcctcatttcagttgaacacaATGGTCTCCAAGGGAGAGGAACTCTTCACCGGAGTCG-<br>TCCCAATCCTCGTCGAGCTCGACGGAGACGTCAACGGACACAAGTTCTCCGTCTCAGGAGAGG-<br>GAGAGGGAGACGCCACCTACGGAAAGCTCACCCTCAAGTTCATCTGCACCACCGGAAAGCTCC-<br>CAGTCCCATGGCCAACCCTCGTCACCACCTTCACTTACGGAGTCCAATGCTTCTCCCGTTACCCA-<br>GACCACATGAAGCGTCACGACTTCTTCAAGTCCGCCATGCCAGAGGGATACGTCCAAGAGCG-<br>TACCATCTTCTTCAAGgtaagtttaaacattaattaatactaactaaccctgattatttaaattttcagGACGACGGA-<br>AACTACAAGACCCGTGCCGAGGTCAAGTTCGAGGGAGACACCCTCGTCAACCGTATCGAGCTCA-<br>AGgtaagtttaaacagttcggtactaactaaccatacatatttaaattttcagGGAATCGACTTCAAGGAGGACG-<br>GAAACATCCTCGGACACAAGCTCGAATACAACTACAACTCCCACAACGTCTACATCATGGCCGA-<br>CAAGCAAAAGAACGGAATCAAGGTCAACTTCAAGgtaagtttaaacatgattttactaactaactaatct-<br>gatttaaattttcagATCCGTCACAACATCGAGGACGGATCTGTCCAACTCGCCGACCACTACCAACA-<br>AAACACCCCAATCGGAGACGGACCAGTCCTCCTCCCAGACAACCACTACCTCTCCACCCAATCC-<br>GCCCTCTCCAAGGACCCAAACGAGAAGCGTGACCACATGGTCCTCAAGGAGTTCGTCACCGC-<br>TGCCGGAATCACCCACGGAATGGACGAGCTCTACATGCCTAAAGATCCAGCCAAACCTCCGGC-<br>CAAGGCACAAGTTGTGGGATGGCCACCGGTGAGATCATACCGGAAGAACGTGATGGTTTCCT-<br>GCCAAAAATCAAGCGGTGGCCCGGAGGCGGCGGCGTTCGTGAAGGTTGAGCAGAAAGAGCA-<br>AGAGCCTATTGTCAA |
| Genotyping primers | |
| Forward | ATCATCACCCATCTCCAACC |
| Reverse | CGCGCATCTTGACAATAGGC |

**Table S3 – Plasmid list**

| Plasmid | Base plasmid | Insert | Used donor modules (SapTrap) |
|---|---|---|---|
| pSMR10 | pMLS257 (Addgene #73716) | *Pvha-6::gfp::bbln-1::N--taglinker::tbb-2_3'UTR* | pSMR13; *C. elegans* optimized GFP; *bbln-1* gBlock; pMLS288 (Addgene #73716); pSMR18 |
| pSMR13 | pHSG298 (Takara Biosciences) | *vha-6* promoter | |
| pSMR18 | pHSG298 (Takara Biosciences) | *tbb-2* 3'UTR | |
| pSMR30 | pMLS257 (Addgene #73716) | *Pvha-6::gfp::BBLN (Hs)::N-taglinker::tbb--2_3'UTR* | pSMR13; *C. elegans* optimized GFP; *BBLN (Hs)* gBlock; pMLS288 (Addgene #73716); pSMR18 |
| pSMR33 | L4440 (Addgene #1654) | *act-5* 3'UTR | |
| pSMR34 | L4440 (Addgene #1654) | *ifp-1* | |
| pSMR35 | L4440 (Addgene #1654) | *ifd-1* | |
| pSMR36 | L4440 (Addgene #1654) | *ifo-1* | |

**Supplemental methods – Sequence data processing**

**Data acquisition and data pre-processing**
To prepare the raw sequence data (provided in FASTQ format) for variant analysis, we first removed adapters and low quality leading or trailing bases using Trimmomatic version 0.38 (Bolger et al., 2014): *java -jar trimmomatic-0.35.jar PE input_forward.fq.gz input_reverse.fq.gz output_forward_paired.fq.gz output_forward_unpaired.fq.gz output_reverse_paired.fq.gz output_reverse_unpaired.fq.gz ILLUMINACLIP:TruSeq3-PE-2.fa:2:30:10 LEADING:3 TRAILING:3 SLIDINGWINDOW:4:15 MINLEN:36*.

Next, reads were aligned to the *C. elegans* reference genome (UCSC genome release ce11) using the Burrows-Wheeler Aligner version 0.7.17 BWA-MEM algorithm (http://bio-bwa.sourceforge.net): *bwa mem -M ce11.fa read1.fq read2.fq > output.sam*.

Third, non-aligned pairs were removed using the sam_bitwise_flag_filter.py script from the Galaxy project (https://galaxyproject.org): *python sam_bitwise_flag_filter.py -f input.sam --flag_column 2 '--0x0001=1' '--0x0002=1' > output.sam*.

Fourth, Picard tools version 2.18.14 (http://broadinstitute.github.io/picard/) and Samtools version 1.9 (http://www.htslib.org/) were used to convert the Sequence Alignment Map (SAM) file into a variant analysis-ready Binary Alignment Map (BAM). A sequence dictionary file was created for the *C. elegans* reference genome (*java -jar picard.jar CreateSequenceDictionary -O ce11.dict -R ce11.fa*) and added to the SAM file (*cat ce11.dict aligned_reads.sam > output.sam*). The SAM file was then sorted by coordinate (*java -jar picard.jar SortSam -I input.sam -O output.sam -SO coordinate*), read group names were added and the file converted to BAM format (*java -jar picard.jar AddOrReplaceReadGroups -I input.sam -O output.bam -RGID rgid -RGLB lib1 -RGPL illumina -RGPU unit1 -RGSM sample_name -SORT_ORDER coordinate*).

Finally, duplicate reads were marked (*java -jar picard.jar MarkDuplicates -I input.bam -O output.bam -M metrics.txt -REMOVE_DUPLICATES true -ASSUME_SORT_ORDER coordinate*) and the BAM file was indexed (*bamtools index -in input.bam*).

**Variant Discovery**
Variant discovery was performed using the Genome Analysis Toolkit (GATK version 4.0.9.0) (https://gatk.broadinstitute.org). First, variants were called per-sample using HaplotypeCaller in genomic variant call format (gVCF) mode: *gatk HaplotypeCaller -R ce11.fa -I input.bam O output.g.vcf.gz -ERC GVCF*. Next, per-sample gVCF files (mutant, heterozygous, and wild-type pools for both mutants) were combined into a multi-sample gVCF file: *gatk CombineGVCFs -R ce11.fasta -V input1.g.vcf.gz -V input2.g.vcf.gz (etc) -O output.g.vcf.gz*. Finally, joint genotyping was performed on the combined gVCF file: *gatk GenotypeGVCFs -R ce11.fasta -V input.g.vcf.gz -O output.vcf*.

**Identification of candidate causative mutations**
To identify candidate causative mutations, we first added genomic variant annotations and functional effect predictions to the VCF file using SnpEff (Version 4.3T with annotation file WBcel235.86) (Cingolani et al., 2012): *java -jar snpEff.jar WBcel235.86 input.vcf > output.vcf*. The relevant data was then exported to a tab-delimited table using GATK: *gatk VariantsToTable -R ce11.fa -F CHROM -F POS -GF AD -F ANN -O output.tab*. This exports the chromosome, position, allelic depth (the number of reads corresponding to the reference and that are variant), and the annotation.

Final analysis was done in Microsoft Excel, by selecting variants present at a frequency >0.8 in the homozygous mutant sample, <0.8 in the heterozygous sample, and <0.1 in the homozygous wild type sample. For both mutant alleles, only a single variant from the resulting selection was predicted to affect a protein coding sequence, which was selected for further analysis.
